# Microfibril-associated glycoprotein 4 forms octamers that mediate interactions with elastogenic proteins and cells

**DOI:** 10.1101/2023.09.22.558839

**Authors:** Michael R. Wozny, Valentin Nelea, Iram Fatima S. Siddiqui, Shaynah Wanga, Vivian de Waard, Mike Strauss, Dieter P. Reinhardt

## Abstract

Microfibrillar-associated protein 4 (MFAP4) is a 36-kDa extracellular glycoprotein with critical roles in human pathologies, including fibrosis in several organ systems, chronic obstructive pulmonary disease, and cardiovascular disorders. In elastic tissues such as arteries, lungs, and skin, MFAP4 associates with microfibrils and elastic fibres, which are the central extracellular fibres affected in thoracic aneurysms. MFAP4 directly interacts with elastogenic proteins, including fibrillin-1 and tropoelastin, and with cells via integrins. MFAP4 multimerisation represents a critical hallmark required for its physiological and pathological properties. However, molecular details and functional consequences of MFAP4 multimerisation are lacking.

Here we present a cryo-electron microscopy structure of human MFAP4. In the presence of calcium, MFAP4 assembles as an octamer with D2 point group symmetry, where two sets of homodimers constitute the top and bottom halves of each octamer. Each of the homodimers is linked together by an inter-molecular disulfide bond. An engineered C34S missense mutation in MFAP4 prevented disulfide-bond formation between monomers, but the mutant formed octamers similar to wild type MFAP4. The atomic model, built into the 3.55 Å cryo-EM map, suggests that several salt-bridges are important for interactions within and between homodimers, while non-polar interactions are important for octamer halves to assemble. In the absence of calcium, MFAP4 dissociates into tetramers, representing the top/bottom halves of the octamers. Binding studies with elastogenic proteins, including fibrillin-1, tropoelastin, LTBP4, and small fibulins showed that MFAP4 has multiple surfaces for protein-protein interactions, which depend upon the higher-order assembly of MFAP4. While the disulfide-bond mediated by C34S contributes little to those protein interactions, it modulated cell interaction. When MFAP4 forms assemblies with fibrillin-1, it abrogates MFAP4 interactions with cells. Overall, the study provides detailed molecular structure-function relationships of MFAP4 interactions with elastogenic proteins and cells.

## Introduction

Microfibril-associated glycoprotein 4 (MFAP4) is a 36 kDa extracellular matrix glycoprotein present in elastic fibre-rich tissues ^1,2^. MFAP4 consists of an N-terminal signal peptide, an RGD recognition site for cell surface integrin receptors, and a C-terminal fibrinogen-related domain (FReD) which constitutes the bulk of the protein (Fig. 1A). MFAP4 partakes in elastin fibre formation (elastogenesis) and in extracellular matrix organisation by protein-protein and cell-matrix interactions. Diseases associated with MFAP4 include Marfan syndrome, cardiovascular disorders, chronic obstructive pulmonary disease, liver fibrosis and cirrhosis, Smith-Magenis syndrome, asthma, and cancer (for review see ^3^).

**Figure 1.**
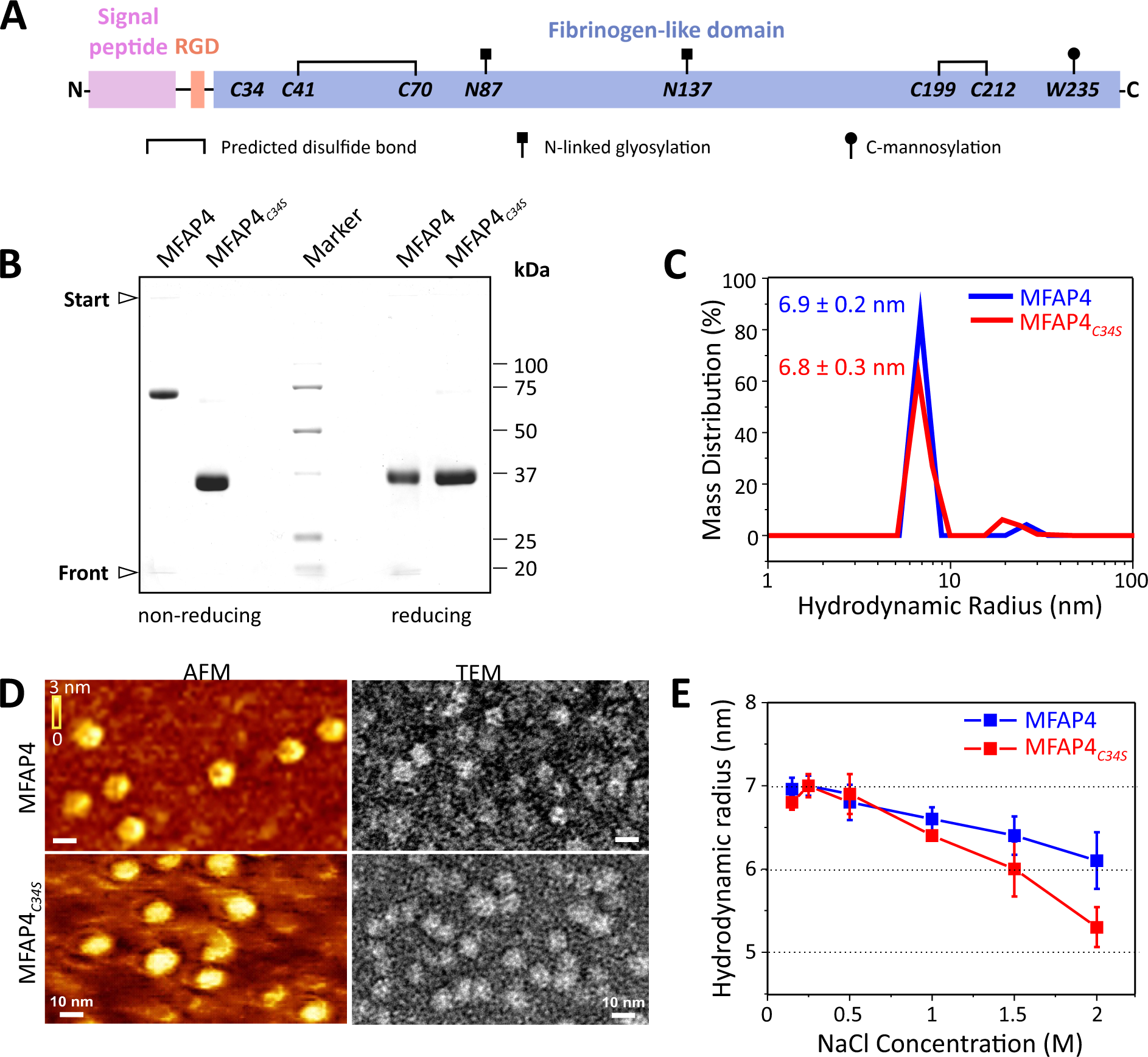
Intermolecular disulfide bonds form between C34 of MFAP4 but are not necessary for multimerisation. (**A**) Schematic of MFAP4 sequence depicting the N-terminal signal peptide, cell surface integrin-binding RGD motif and the fibrinogen-related domain (FReD) of human MFAP4. Cysteine residues, N-glycosylation and C-mannosylation sites are indicated. In MFAP4*_C34S_*, cysteine 34 is altered to serine. Schematic adopted from Kanaan *et al.* ^3^. (**B**) Coomassie-stained gels after SDS-PAGE of MFAP4 and MFAP4*_C34S_* under non-reducing (left lanes) and reducing (right lanes) conditions. (**C**) Hydrodynamic radii of MFAP4 (blue) and MFAP4*_C34S_* (red) in TBS/Ca^2+^, measured by DLS. (**D**) AFM and negative stain TEM of MFAP4 (top panels) and MFAP4*_C34S_* (bottom panels). Scale bars are 10 nm for all images. (**E**) NaCl-dependent hydrodynamic radii of MFAP4 (blue) and MFAP4*_C34S_* (red), determined by DLS.

Elastogenesis represents a highly hierarchical universal process that starts with fibronectin network formation which is strictly required for the formation of “bead-on-the-string” fibrillin-1 containing microfibrils ^4-7^. These microfibrils provide a cell surface-associated scaffold for the deposition of tropoelastin as nanofibres, which possibly interact with cells via integrins and heparan sulfate ^8,9^. Several accessory proteins play crucial roles in elastic fibre formation and homeostasis, including MFAP4, fibulin-4, fibulin-5, latent transforming growth factor beta binding protein-4 (LTBP4), and lysyl oxidases, among others ^10-16^. Mature elastic fibres are comprised of an inner core of crosslinked elastin, an outer mantle of microfibrils and an interface connecting both ^4,17^.

A series of studies have substantiated the role of MFAP4 in elastogenesis. It is present at the elastic fibre-microfibril interface ^1,18^, and it typically associates with fibrillin-containing microfibrils in many elastic tissues including aorta, lung, and skin ^2,10,18-20^, but not with elastin-free microfibrils in the ciliary zonules of the eye or the kidney mesangium ^18^. Adult mice with a genetic deletion of *Mfap4* show impaired lung elastic properties and emphysema-like alterations ^21^, and carotid artery ligation delayed neointima formation ^22^. Protein expression levels are elevated in several diseases including chronic obstructive pulmonary disease ^23^, abdominal aortic aneurysms ^24^, or hepatic fibrosis ^25^. MFAP4 is upregulated in the extracellular matrix of the ascending aortae from individuals with Marfan syndrome and high circulating plasma MFAP4 levels were associated with type B aortic dissections in the descending thoracic aorta ^26^. Glycoproteomic analysis identified increased and more diverse N-glycosylation patterns of MFAP4 in patients with Marfan syndrome compared to controls ^26^.

MFAP4 forms a disulfide-bonded dimer, that further assembles into higher oligomers ^27^. However, the exact composition and identity of those oligomers remains controversial. Based on biochemical studies, Schlosser *et al.* reported MFAP4 tetramers of homodimers (i.e. octamers) ^27^, whereas Pilecki *et al.* determined trimeric and hexameric structures of homodimers (i.e. hexamers and dodecamers, respectively) ^10^. Irrespective of the multimeric structure, the prevailing paradigm is that disulfide-bonded monomers represent the core assembly unit. Multimerisation of other elastogenic proteins has significant functional consequences, for example as described for fibrillin- 1 and -2 ^7^, as well as for fibulin-4 and LTBP-4 ^11,28^. Thus, it is predicted that the multimeric state of MFAP4 determines its specific functions. Whether N-linked glycosylation at position 87 and 137 or mannosylation at position 235 play a role in MFAP4 oligomerisation and function remains to be established ^26,29^. With respect to elastogenesis, MFAP4 directly interacts with tropoelastin and the elastin-specific cross-link desmosine in a calcium-dependent fashion, and with fibrillin-1 and -2 as well as with lysyl oxidase in a calcium-independent manner ^10,30^. In addition, MFAP4 promotes coacervation of tropoelastin ^10^, and interacts with elastogenic cells via an RGD motif that interacts with integrin αvβ3 and possibly αvβ5, thereby promoting cell migration and proliferation ^22,30^. Based on homology with related FReD family proteins as well as on the analysis by the AlphaFold software, it is predicted that four of the five cysteines within MFAP4 form intramolecular disulfide bonds (Fig. 1A) ^3,31,32^, which leaves the possibility that the fifth N-terminal C34 residue participates in an intermolecular disulfide bond between MFAP4 monomers. However, this prediction requires experimental validation, and the relationship between MFAP4 oligomerisation and its specific functions is virtually unknown.

Despite important advances in understanding the structural and functional relationships of elastogenic proteins in general, and MFAP4 in particular, critical molecular mechanisms that drive elastic fibre assembly remain unclear without sub-nanometer structural information. Here we make use of single particle analysis (SPA) with cryogenic transmission electron microscopy (cryo-EM) to investigate the MFAP4 ultrastructure, determine its atomic model structure and assess its macromolecular assembly. We combine this novel information with experiments to reveal structure-function relationships with respect to elastogenic mechanisms.

## Results

### Multimeric organisation of MFAP4 is not dependent upon intermolecular disulfide-bond formation

To investigate the oligomeric state of MFAP4, and whether C34 is necessary for intermolecular disulfide-bond formation within a dimer, we produced highly purified recombinant human MFAP4 and MFAP4*_C34S_* using HEK293 cells. As described previously by others ^10^, MFAP4 migrated in SDS gels as a ∼72 kDa band (dimer) under non-reducing conditions and as a ∼36 kDa band (monomer) under reducing conditions (Fig. 1B). In contrast, MFAP4*_C34S_* migrated as a ∼36 kDa band under both non-reducing and reducing conditions (Fig. 1B). This data provides evidence that C34 is required and sufficient for intermolecular disulfide-bond formation between monomers to stabilise dimers. In solution under physiological buffer conditions (TBS/Ca^2+^), dynamic light scattering (DLS) of both MFAP4 and MFAP4*_C34S_* demonstrated a similar hydrodynamic radius of 6.9 ± 0.2 nm and 6.8 ± 0.3 nm, respectively (Fig. 1C). Based on DLS particle size calibration with standard proteins, this is consistent with a molecular mass of approximately 280 kDa, suggesting that both MFAP4 and MFAP4*_C34S_* assemble as a complex of eight monomers in the presence of Ca^2+^ (Supp. Fig. S1A). Similarly, MFAP4 in solution dissociated into dimers upon treatment with 1% SDS, causing a shift in hydrodynamic radius to 3.7 ± 0.2 nm (Supp. Fig. S1B). When MFAP4 was treated with 50 mM reducing dithiothreitol (DTT) and 1% SDS, dimers were reduced to monomers with a hydrodynamic radius of 2.7 ± 0.1 nm (Supp. Fig. S1B). Irrespective of DTT treatment, MFAP4*_C34S_* dissociated into monomers in the presence of 1% SDS with a hydrodynamic radius of 2.6 ± 0.2 nm (Supp. Fig. S1B). Relatively low levels of DTT (∼3 mM) were required for 50% reduction of MFAP4 and conversion of dimers to monomers, suggesting a surface-located position for the intermolecular C34 disulfide bond (Supp. Fig. S1C). When analysed by atomic force microscopy (AFM) or negative-stain transmission electron microscopy (TEM), both MFAP4 and MFAP4*_C34S_* appeared as round particles ∼15 nm in diameter by AFM and ∼12 nm in diameter by TEM (Fig. 1D); again, indicative of higher-order oligomeric assemblies. DLS analysis of MFAP4 under high salt conditions (2 M NaCl at a physiological pH 7.4) reduced the hydrodynamic radius of particles to 6.2 ± 0.3 nm (Fig. 1E). Interestingly, increasing NaCl concentrations affected the hydrodynamic radius of MFAP4*_C34S_* more strongly than that of MFAP4 (5.3 ± 0.2 nm at 2 M NaCl) (Fig. 1E), suggesting that C34 stabilises MFAP4 assemblies but is not necessary for the formation of higher ordered assemblies.

### MFAP4 dimers assemble to form an octamer

SPA of MFAP4 cryo-EM data revealed similar structures to those observed by AFM and negative-stain TEM (Fig. 2A). Initial 2D class averages consisted of two predominant views of four lobes; one where the four lobes are equidistant to each other with a ellipsoidal cavity between these lobes and a second arrangement in which the cavity between the lobes is elongated along a single axis (Fig. 2B). The cavity opening dimensions are ∼1.9 nm wide and ∼3.5 nm high. These two views are consistent with top/bottom *vs* side views of an octamer which is elongated along one axis when viewed from its side. Furthermore, an octameric arrangement of MFAP4 was suggested by 2D class averages which appeared to be projections of eight globular MFAP4 molecules each perched at the vertex of a cube. The presented 2D classes were collectively used for initial reference generation and subsequent 3D reconstruction using C1 point group symmetry, confirming that MFAP4 organises as an octameric structure. Further refinement of the 3D cryo-EM map with D2 symmetry produced a 3.55 Å resolution (masked FSC*_half-map_ =* 0.143, Supp. Fig. S2) structure of octameric MFAP4 (Fig. 2C, Movie 1). This cuboid assembly was elongated along the axis drawn between the top and bottom faces of this complex. Despite the inner cavity, the individual components appeared closely packed.

**Figure 2.**
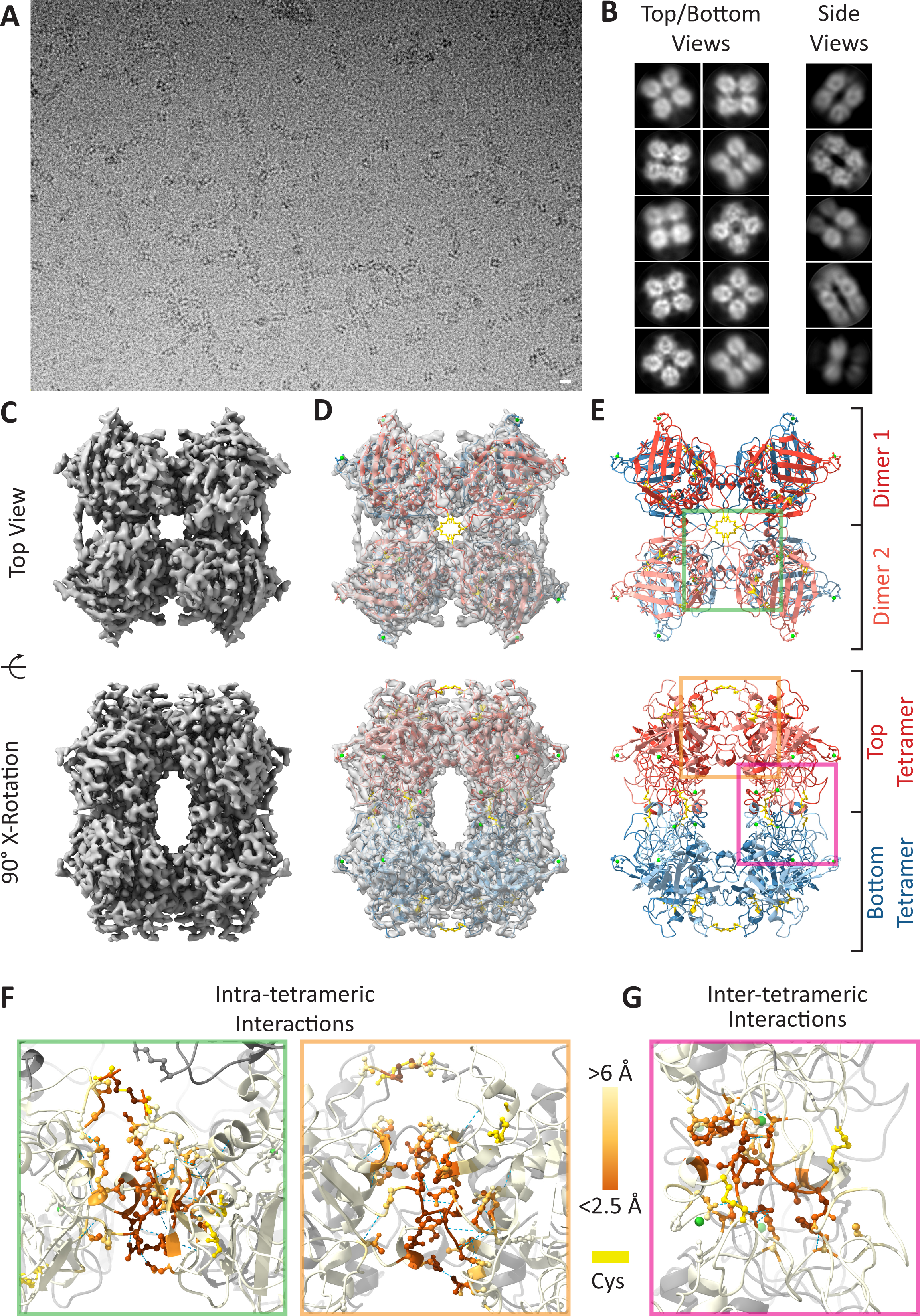
SPA of MFAP4 with Ca^2+^. (**A**) Cryo-EM micrograph of MFAP4 with Ca^2+^ (scale bar 10 nm). (**B**) 2D-class averages of top/bottom and side views of MFAP4 particles used for reconstruction of initial 3D reference. (**C**) Top and side views of the D2 symmetry 3.55 Å resolution 3D cryo-EM map of MFAP4, (**D**) along with superimposed atomic model, and (**E**) the atomic model. The top tetramer is red with the dimer halves coloured light and dark shades of red. The bottom tetramer is coloured blue with dimer halves coloured light and dark shades of blue. (**F**) Intra-tetrameric interactions as seen from the top (green inset) and side view (orange inset) of subpanel E. (**G**) Inter-tetrameric interactions as seen from the side view (pink inset) of subpanel E. Cysteine residues are coloured yellow in all subpanels. Hydrogen bonds are depicted as dashed blue lines. Residues are coloured according to inter-chain distance (dark orange <2.5 Å, light orange <8.5 Å; see colour scale in **F/G**).

An AlphaFold2-predicted model of MFAP4 was positioned within this cryo-EM map and further refined through iterative manual and automatic adjustment of atomic coordinates using Coot ^33^ and Phenix ^34^ (Fig. 2D; Movie 1). With this map, we could accurately resolve the positions of residues Q37-A255 (fixed-radius masked crossed correlation score = 0.84). We observed cryo-EM map density compatible with intramolecular disulfide bonds between the modelled positions of C41-C70 and C199-C212. Although the N-termini preceding Q37 are not resolved within the map-model and are predicted to be disordered regions by AlphaFold2, we observed that the Q37 residues of each of the four MFAP4 molecules at the top and bottom of an octamer are positioned to extend towards each other and outwards from the macromolecular assembly (Movie 1). This arrangement potentially positions four integrin binding RGD motifs at each of the poles of the MFAP4 octamer (Movie 1, orange residues). Furthermore, we observed that intermolecular disulfide bonds between C34 residues of different MFAP4 monomers are possible for this model when constrained by the cryo-EM map, although we do not resolve these residues within the cryo-EM map. We modelled these intermolecular disulfide bonds to reflect the dimerisation of MFAP4 under non-reducing conditions in SDS gels. In agreement with the disruption of MFAP4 dimers by low concentrations of dithiothreitol (Fig. 1B and Supp. Fig. S1C), these disulfide bonds are exposed to the solvent and are readily available for a reducing agent. Thus, the top and bottom halves of the MFAP4 octamer, defined across a plane perpendicular to the octamer’s longest axis. These halves can be described as two identical tetramers each composed of two dimers with N-termini linked by C34 disulfide bonds.

Within the tetrameric top and bottom halves of the MFAP4 octamer, monomers interact at multiple contact points through intra-tetrameric interfaces (Fig. 2E,F, Movie 2), including potential salt-bridge interactions between interchain residues R105-D159 and R94-E158, hydrogen bonds between interchain residues A106-N152, G104-E158, G104-V154 and R94-E158, as well as non-bonded interchain interactions between R105-N152, G104-D159 and F103-E158. These interactions occur between each monomer so that there are four interaction surfaces comprised of salt-bridge, hydrogen bonds and non-bonded interchain interactions per tetrameric half of an MFAP4 octamer. The buried solvent-accessible surface area – a measure of the extent of the interaction surface, or area not accessible to solvent – of these intra-tetrameric interaction surfaces is 615 or 633 Å^2^, depending on whether the monomers are disulfide bond linked. Each interface is shared between 40 contacting residues (16 residues of one chain interact with 24 residues of another chain). The top and bottom halves of the MFAP4 octamer associate through inter-tetrameric interfaces (Fig. 2G), including a polar non-bonded interchain interaction between N231-A200 and non-polar non-bonded interchain interactions between V196-F93, L202-L202, S204-L202 (Fig. 2G, Movie 3). Between tetrameric halves of the MFAP4 octamer, the buried solvent-accessible surface area is 433Å^2^ and is shared between 22 contacting residues (13 residues from each adjacent chain).

Noteworthy, these interacting interfaces between tetramers correspond to regions within the previously described S1 binding site of MFAP4 (T187-K251) ^10^. Previously resolved structures of other FReD proteins show that this site contains a Ca^2+^-binding site. Ca^2+^-autoradiography demonstrated that bovine MAGP-36 binds calcium ^35^; however, one or more definitive Ca^2+^-binding sites have not been resolved as alterations of potential Ca^2+^-binding residues disrupted protein stability and expression ^10^. We used ConSurf ^36^ to identify other conserved functional regions within the MFAP4 model (Supp. Fig. 3). Upon close inspection of loop A182-A206 as a candidate for Ca^2+^-binding, we noticed cryo-EM map density between S186, T187, D191 and Q192 that was not explained by the atomic model of MFAP4. D191 and D193 of MFAP4 are conserved across several FreD proteins ^10^, and equivalent residues in Fibrinogen C Domain Containing 1 protein (FIBCD1) bind a Ca^2+^ ion ^31^. To account for this, we positioned a Ca^2+^ ion within the cryo-EM map density near D191. We noticed that S186, D191 and Q192 sidechains as well as the backbone carbonyl group of T187 were all capable of binding Ca^2+^ at this position. We propose that this Ca^2+^ helps to push loop A182-A206 away from MFAP4’s centre of mass and towards the opposing A182-A206 loop of the other MFAP4 molecules, thus stabilising inter-tetrameric interactions between V196-F93, L202-L202, S204-L202 and N231-A200 to form octamers. We note that another conserved residue, D134, was observed to form a similar Ca^2+^-binding pocket with E136 and N138. These residues are close to N137, which has previously been reported to be glycosylated ^37^, and thus may affect the stability of a glycan attached to N137.

### Structural evidence of glycosylation in MFAP4

Cryo-EM map densities reach out from the MFAP4 octamer at the positions N87 and N137, supporting that both sites are N-glycosylated (Fig. S4A, B; Movie 1, blue residues). We modelled a N-acetyl-D-glucosamine (NAG), the first unit of N-linked oligosaccharides, within the cryo-EM density nearby to both N87 and N137. Although the cryo-EM density at these positions extends beyond the modelled NAG molecules, as is common for variable glycosylation patterns, we were unable to fit further monosaccharides reliably in this extended domain. Only the first monosaccharide of the glycan at N137 can be discerned above the noise within the cryo-EM map, while the glycan at N87 is discernible for at least two of its constituent monosaccharides; although we only model one NAG at both N87 and N137. The cryo-EM density near N87 is visible at a higher contour than that around N137, suggesting that this glycan is less flexible or more common. The N87 glycan is positioned near the central cavity formed between the two halves of the MFAP4 octamer, while the N137 glycan is positioned at the periphery of the octamer surface. W235 was reported to be mannosylated, and in the absence of this modification MFAP4 secretion was decreased ^29^. However, there is no cryo-EM map density around W235 that could correspond to a glycan (Supp. Fig. S4C). W235 is buried within MFAP4 and not accessible as a surface residue. As such, W235 appears to play a structural role within the MFAP4 monomer and forms hydrogen bonds with A173, W238, and K239 through backbone-backbone interactions.

### MFAP4 octamers disassemble to tetramers when calcium is depleted

DLS measurements of MFAP4 and MFAP4*_C34S_* showed that hydrodynamic radii decreased from 6.9 ± 0.2 nm and 6.8 ± 0.3 nm in the Ca^2+^-loaded form (Fig. 1C) to 5.3 ± 0.3 nm and 5.2 ± 0.4 nm in the Ca^2+^-depleted form (Fig. 3A). These values are consistent with the possibility that ∼280 kDa MFAP4 octamers in the Ca^2+^ form dissociate into particles approximately half the molecular mass (∼140 kDa) when Ca^2+^ is removed (see calibration curve in Supp. Fig. S1A). Interestingly, surface plasmon resonance spectroscopy (SPR) showed that MFAP4 octamers self-interact with a K*_D_* = 6.2 ± 0.9 nM in the presence of Ca^2+^, whereas only negligible self-interaction occurred upon removal of Ca^2+^ (Fig. 3B). This data suggests that the conformation of the MFAP4 octamer is important for self-interaction. Similar self-interaction properties were observed for MFAP4*_C34S_* (K*_D_* = 5.8 ± 1.5 nM in the Ca^2+^ form, negligible interaction in the Ca^2+^-depleted form) (Fig. 3C), demonstrating that C34-mediated intermolecular disulfide bonds are neither required for self-interaction nor for the Ca^2+^-mediated assembly of the MFAP4 octameric structure.

**Figure 3.**
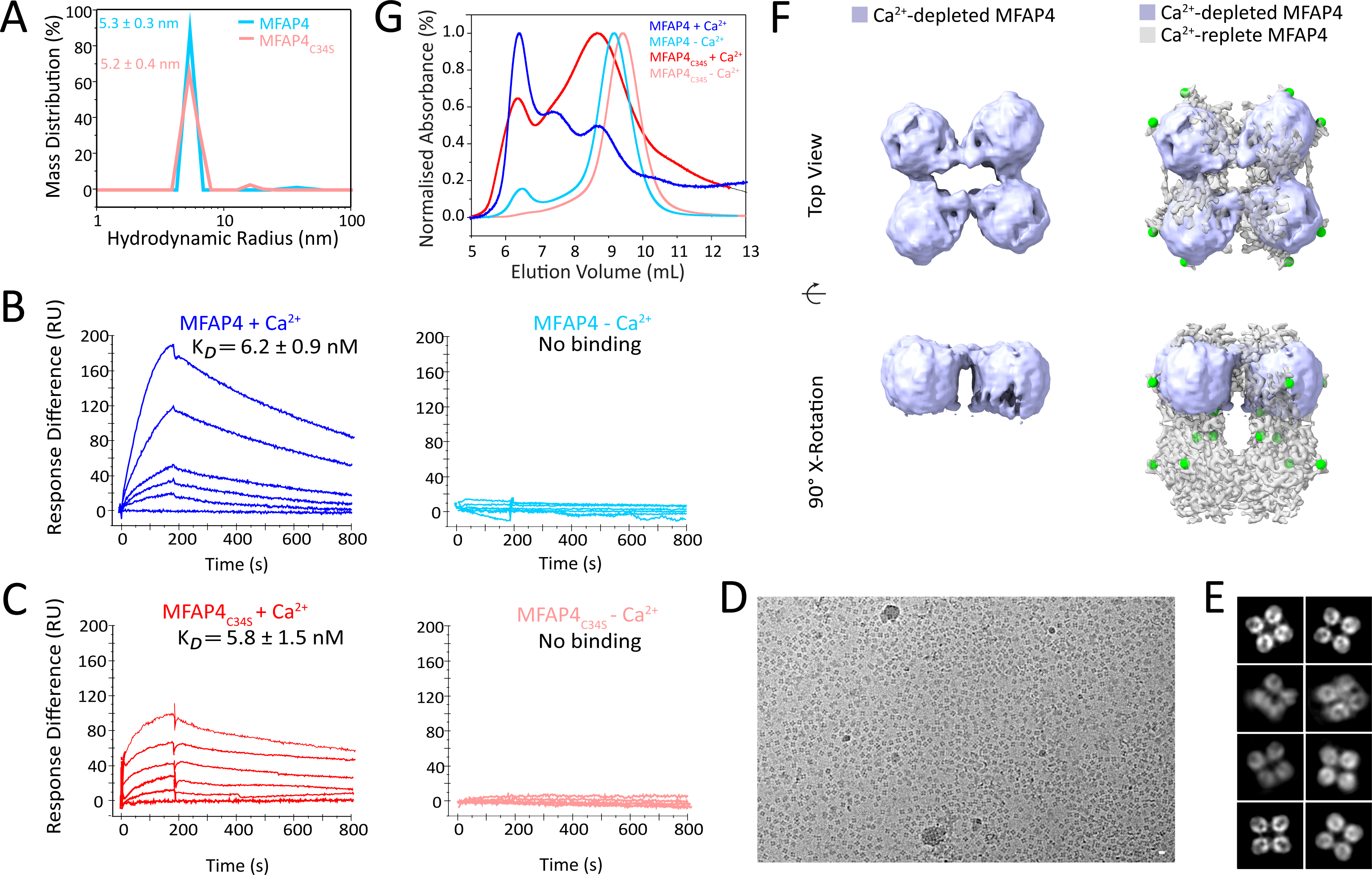
Ca^2+^ is necessary for MFAP4 octamer formation and self-interactions. (**A**) Hydrodynamic radii of MFAP4 and MFAP4*_C34S_* in TBS without Ca^2+^ after treatment with EDTA, measured by DLS. (**B**) Self-interaction of MFAP4 (**B**) or MFAP4*_C34S_* (**C**) measured by SPR with (+Ca^2+^) or without (-Ca^2+^; EDTA treated) calcium. (**D**) Cryo-EM micrograph of MFAP4 in TBS without Ca^2+^ after EDTA treatment (scale bar 10 nm). (**E**) 2D-class averages of top/bottom of EDTA-treated MFAP4. (**F**) C2 3D reconstruction of EDTA-treated MFAP superimposed within the top half of the D4 reconstruction of MFAP4 with Ca^2+^. (**G**) Gel filtration of MFAP4 and MFAP4*_C34S_* in TBS either in the presence (+ Ca^2+^) or absence (- Ca^2+^; EDTA treated) of calcium.

The shift in hydrodynamic radius of both MFAP4 and MFAP4*_C34S_* upon removal of Ca^2+^ suggested a prominent structural change and prompted us to investigate Ca^2+^-depleted MFAP4 by SPA with cryo-EM. Micrographs of EDTA-treated MFAP4 showed structural assemblies similar to those of MFAP4 in the Ca^2+^form; however, the distribution of the particles was more evenly dispersed within the ice (Fig. 3D) as compared to MFAP4 with Ca^2+^ (Fig. 2A). 2D classes of EDTA-treated MFAP4 resembled top/bottom view 2D classes of the MFAP4 octamer in the Ca^2+^- form. However potential side view 2D classes were absent from EDTA-treated MFAP4 particles (Fig. 3E). An asymmetric 3D reconstruction suggested a tetrameric structure. Refinement of the EDTA-treated MFAP4 cryo-EM map proved challenging, as it became clear that proteins adopted a strong preferential orientation during vitrification on cryo-EM support grids (Supp. Fig. S5). The refined tetramer structure of EDTA-treated MFAP4 fits within either top or bottom half of the Ca^2+^-loaded MFAP4 octamer structure (Fig. 3F). We suspect that an increase in flexibility of the tetramer limits the resolution of the reconstruction. To further characterise the oligomeric state of Ca^2+^-depleted MFAP4 biochemically, we used gel filtration chromatography of MFAP4 and MFAP4*_C34S_* in the presence and absence of EDTA. We observed three distinct peaks between 6-10 mL elution volume for the Ca^2^-form of MFAP4 and MFAP4*_C34S_*. When MFAP4 and MFAP4*_C34S_* were treated with EDTA, we noted a strong decrease in the absorbance of elution volumes corresponding to the first two peaks (6-8 mL) and a shift of the third peak (8-10 mL) to smaller particle sizes (Fig. 3G). Together, the data support a model where Ca^2+^ is necessary for the assembly and/or stabilisation of octamers but not of tetramers. We propose that in the absence of Ca^2+^-binding by S186, T187, D191 and Q192, the conformation of loop A182-A206 changes thus preventing inter-tetrameric interactions between V196-F93, L202-L202, S204-L202 and N231-A200 to form octamers. Ca^2+^-binding by D134, E136, and N138 does not appear to be relevant to octamer formation.

### MFAP4 and MFAP4*_C34S_* interaction properties with respect to structural aspects

Since MFAP4 associates with fibrillin-containing microfibrils, and binds to fibrillin-1 and tropoelastin ^10^, we tested interactions with a series of elastogenic proteins and protein fragments using SPR to understand how MFAP4 multimerisation and C34 are required for those interactions (Fig. 4 and Supplemental Table 1). In the presence of Ca^2+^ (octameric form), MFAP4 interacted with the N-terminal half (rFBN1-N, K*_D_* = 1.8 ± 0.9 nM) and with the centre (rF1M, K*_D_* = 1.2 ± 0.5 nM) of fibrillin-1 with very high affinity. Interactions with tropoelastin (K*_D_* = 45 ± 9 nM), LTBP4L (K*_D_* = 18 ± 5 nM), and LTBP4S (K*_D_* = 15 ± 6 nM) were of high affinity, the latter mediated by the N-terminal half of LTBP4L and LTBP4S (Supplemental Table 1). Binding to fibulin-4 (K*_D_* = 110 ± 33 nM) and fibulin-5 (K*_D_* = 357 ± 46 nM) were of lower affinity. Fibronectin, fibulin-3, and the C-terminal half of fibrillin-1 (rFBN1-C) did not interact with the Ca^2+^-loaded MFAP4 (Supplemental Table 1). After Ca^2+^ was removed (tetrameric form), MFAP4 did not interact with the fibrillin-1 fragments, tropoelastin, fibulin-4 or fibulin-5, whereas MFAP4 interaction with LTBP4L and LTBP4S was not affected by Ca^2+^ depletion (Fig. 4). These data are consistent with the interpretation that the fully assembled MFAP4 octamer is required for interaction with fibrillin-1, tropoelastin, fibulin-4, and fibulin-5, whereas the MFAP4 tetramer is sufficient to interact with LTBP4L and LTBP4S. For MFAP4*_C34S_*, relatively similar binding properties were observed in the Ca^2+^ form, except for fibulin-5 that did not interact (Fig. 4). None of the ligands could interact with MFAP4*_C34S_* upon removal of Ca^2+^, including those interactions that occurred with MFAP4 in the Ca^2+^-depleted form (LTBP4L, LTBP4S). These data suggest that the C34-mediated intermolecular disulfide bonds are not necessary for protein interactions requiring octameric MFAP4, but they do affect those interactions that only require the MFAP4 tetramer.

**Figure 4.**
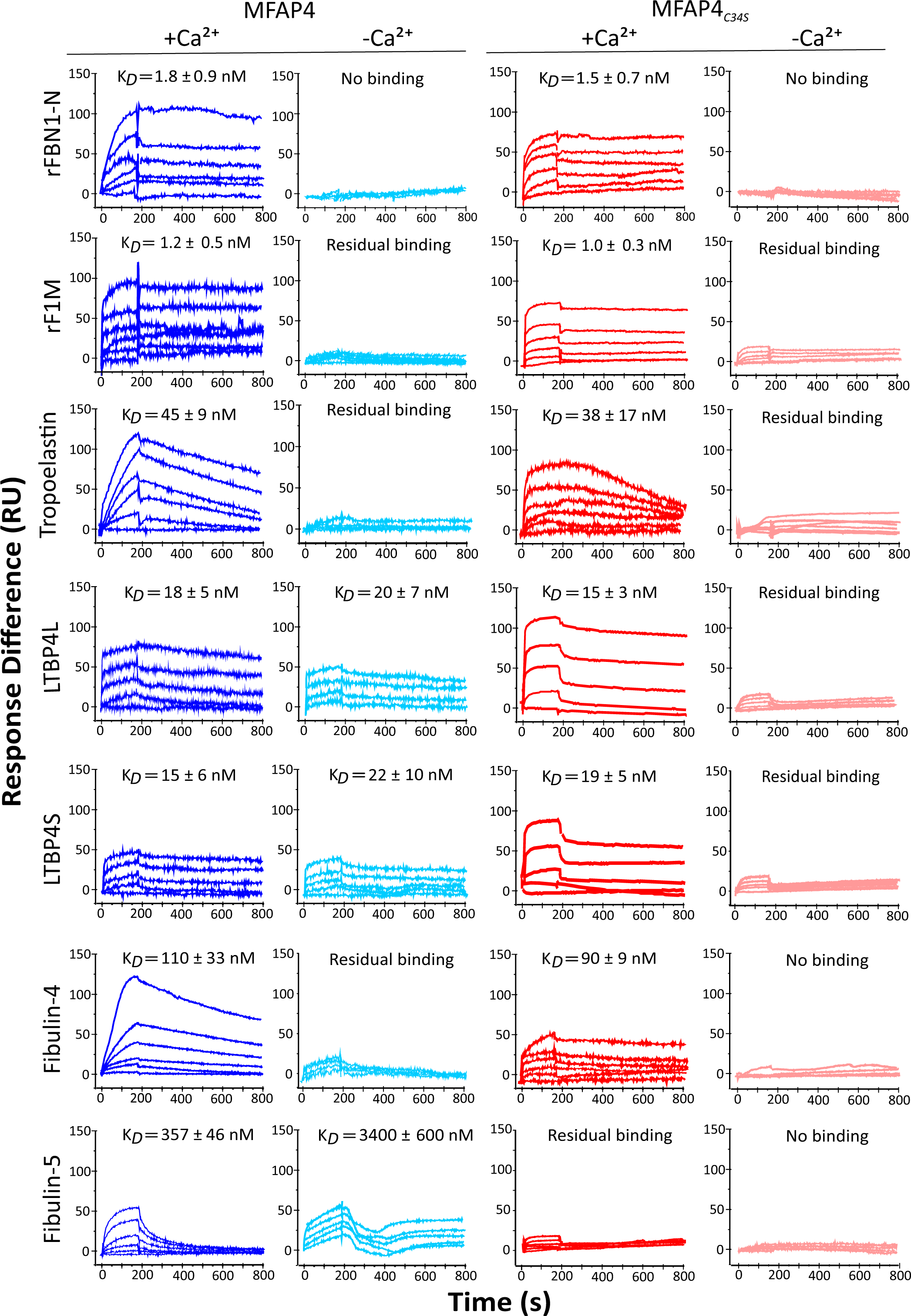
MFAP4 and MFAP4*_C34S_*interactions with elastic fibre-associated extracellular matrix proteins. The interactions were tested by SPR with MFAP4 and MFAP4*_C34S_* immobilized on the sensor surface either in the presence (+Ca^2+^) or absence (-Ca^2+^; EDTA treatment followed by TBS dialysis) of calcium. Full-length proteins and subfragment proteins (as indicated) were used as analytes in the soluble phase (100, 50, 20, 10, 5, 2 and 0 µg/mL for each sensorgram) either in TBS containing Ca^2+^ (+Ca^2+^) or in TBS without calcium (-Ca^2+^). Additional tested interactions are shown in Supplemental Table 1.

Since the RGD motif for cell interaction site is located close to C34, we tested attachment of human dermal fibroblasts to MFAP4 and MFAP4*_C34S_* (Fig. 5A). MFAP4*_C34S_* promoted fibroblast attachment slightly but significantly more than MFAP4. These data suggest that the RGD site within the N-terminus of MFAP4*_C34S_* is more readily available to the respective cell surface integrins, likely due to less constraint than normally provided by the C34-mediated intermolecular disulfide bonds between MFAP4 monomers. Since MFAP4 interacts with fibrillin-1 with very high affinity at the N-terminus, we analysed complexes of the N-terminal half as well as full length recombinant fibrillin-1 with MFAP4 by AFM (Fig. 5B). Particles consistent with the diameter of the MFAP4 octamer occurred frequently at the end of rFBN1-N and full length fibrillin-1. In some instances, longer assemblies of full length fibrillin-1 were observed with an MFAP4 spacing of about one fibrillin-1 molecule length (∼130 nm). This data suggested that MFAP4 interacts with the fibrillin-1 N-terminus close to the site of N-to-C fibrillin-1 self-interactions ^7^. We further analysed whether interactions of MFAP4 and MFAP4*_C34S_* with cells were modulated upon fibrillin-1 binding. When the MFAP4 proteins were either pre-mixed with rFBN1-N before coating the wells, or when they were first coated and then incubated with rFBN1-N, adhesion to fibroblasts were abolished (Fig. 5C). This data shows that the RGD motif at the octamer poles become inaccessible for cell surface integrins upon fibrillin-1 binding, either through a competitive interaction mechanism or through steric hindrance.

**Figure 5.**
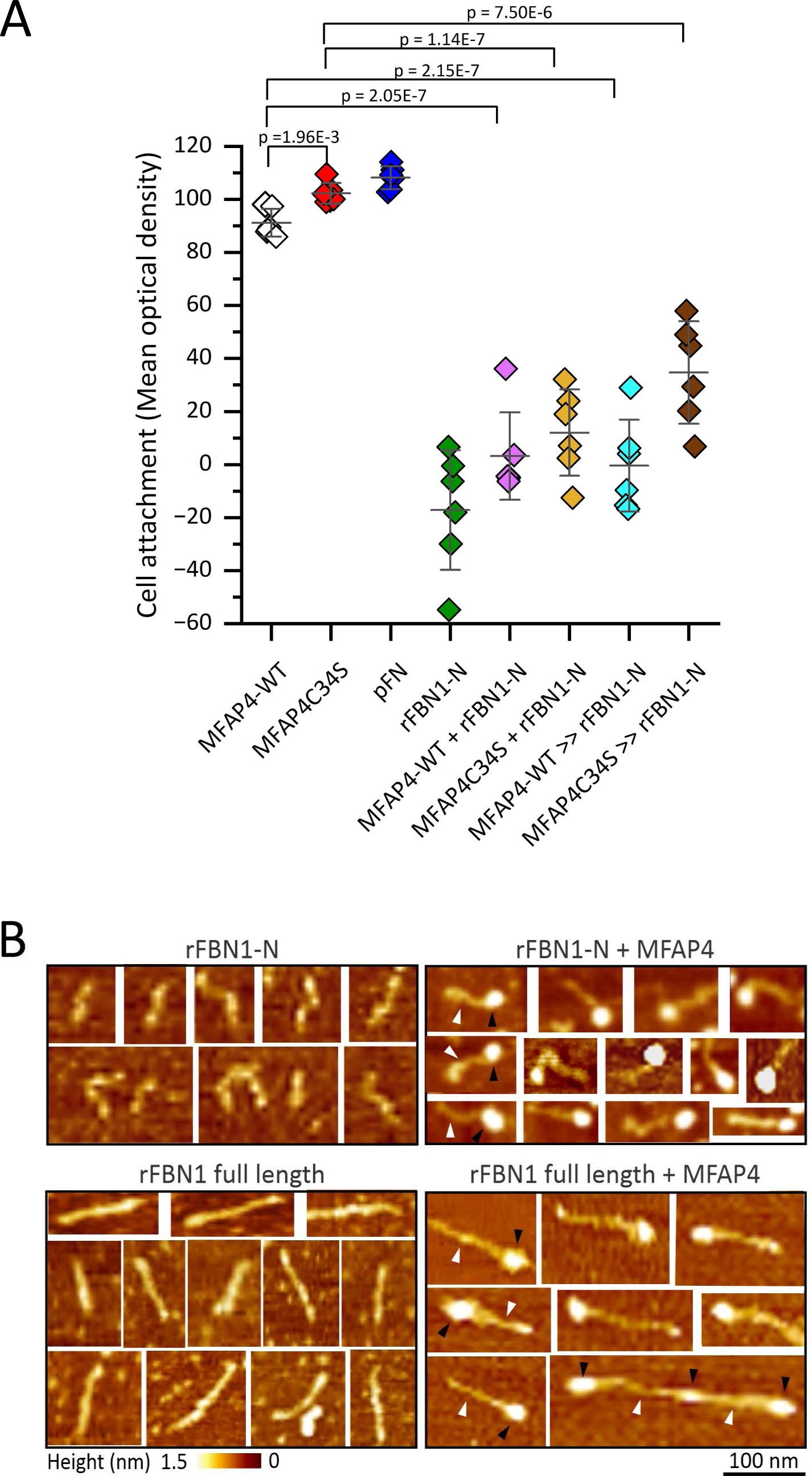
MFAP4 and MFAP4*_C34S_* interactions with fibroblasts and visualisation of MFAP4-fibrillin-1 complexes. (**A**) Cell attachment of human dermal fibroblasts to MFAP4, MFAP4*_C34S_*, rFBN1-N, and plasma fibronectin (pFN) as control. The proteins were either coated alone as indicated or were first mixed (MFAP4/MFAP4*_S34S_* + rFBN1-N) and then coated, or MFAP4/MFAP4*_S34S_* were first coated followed by incubation with rFBN1-N ((MFAP4/MFAP4*_S34S_* >> rFBN1-N). Shown is one representative experiment with six technical replicates (n=6). Background values of wells not coated with proteins but including cells were subtracted from the experimental values. This correction results for some samples in negative values representing zero binding. Statistical analysis was performed using the two-sample t-test; error bars represent standard deviations; p-values are indicated for each comparison. (**B**) AFM visualization of the recombinant N-terminal half of fibrillin-1 (rFBN1-N; top panel) or full-length recombinant fibrillin-1 (rFBN1; bottom panel) either alone (left panels) or mixed with MFAP4 (right panels). Some rFBN1-N and rFBN1 molecules are marked by white arrowheads, and some MFAP4 molecules are labelled with black arrowheads. Distance scale bar represents 100 nm for all images; height scale bar represents 0-1.5 nm for all images.

### MFAP4 forms chains of octamers

In addition to isolated MFAP4 octamers, extended chain-like structures were visible in micrographs of MFAP4 vitrified in TBS/Ca^2+^ (Fig. 2A). We investigated these higher-order assemblies by reanalysing the aligned dataset, extracting particles with a larger box size (∼723 Å box-width) to capture partner octamers and iterative C1 classification and C1 refinement with a spherical mask (520 Å in diameter) which included both the central octamer and the nearby partner. This scheme yielded a cryo-EM density map with a central octamer and a single partner, suggesting that these chain-like assemblies are sufficiently flexible to be structurally disordered beyond two units (∼266 Å). To further investigate the interface between these partners, we utilised multibody refinement within Relion4 to focus two individual maps on either the central octamer or its partner^38^. Multibody refinement yielded two maps with extensions reaching towards each other (Fig. 6A; Movie 4). Principle component analysis shows that the primary and secondary modes of movement consist of the partner swinging +/- 30° relative to the central body (Fig. 6B, red and yellow partners). The tertiary mode of movement consists of the partner octamer twisting around an axis of connection between the two molecules (Fig. 6B, blue partner). We ascribe these extensions, which were not resolved in the D2 map of a single octamer, to the sequence between Q21, the first residue after signal peptide cleavage, and C34, which is involved in the intermolecular disulfide bond within a single octamer. From these observations, we propose a pole-to-pole organisation of MFAP4 octamers interacting with each other through N-termini directed from the top and bottom of each octamer resulting in linear chains of octamers. DLS of MFAP4 reveals that particle size increases with increasing pH (Fig. 6C); possibly reflecting an importance for electrostatic interactions in mediating MFAP4 chain formation. Whether MFAP4 forms chains *in situ* and whether this may be regulated by RGD cell interactions is currently unknown.

**Figure 6.**
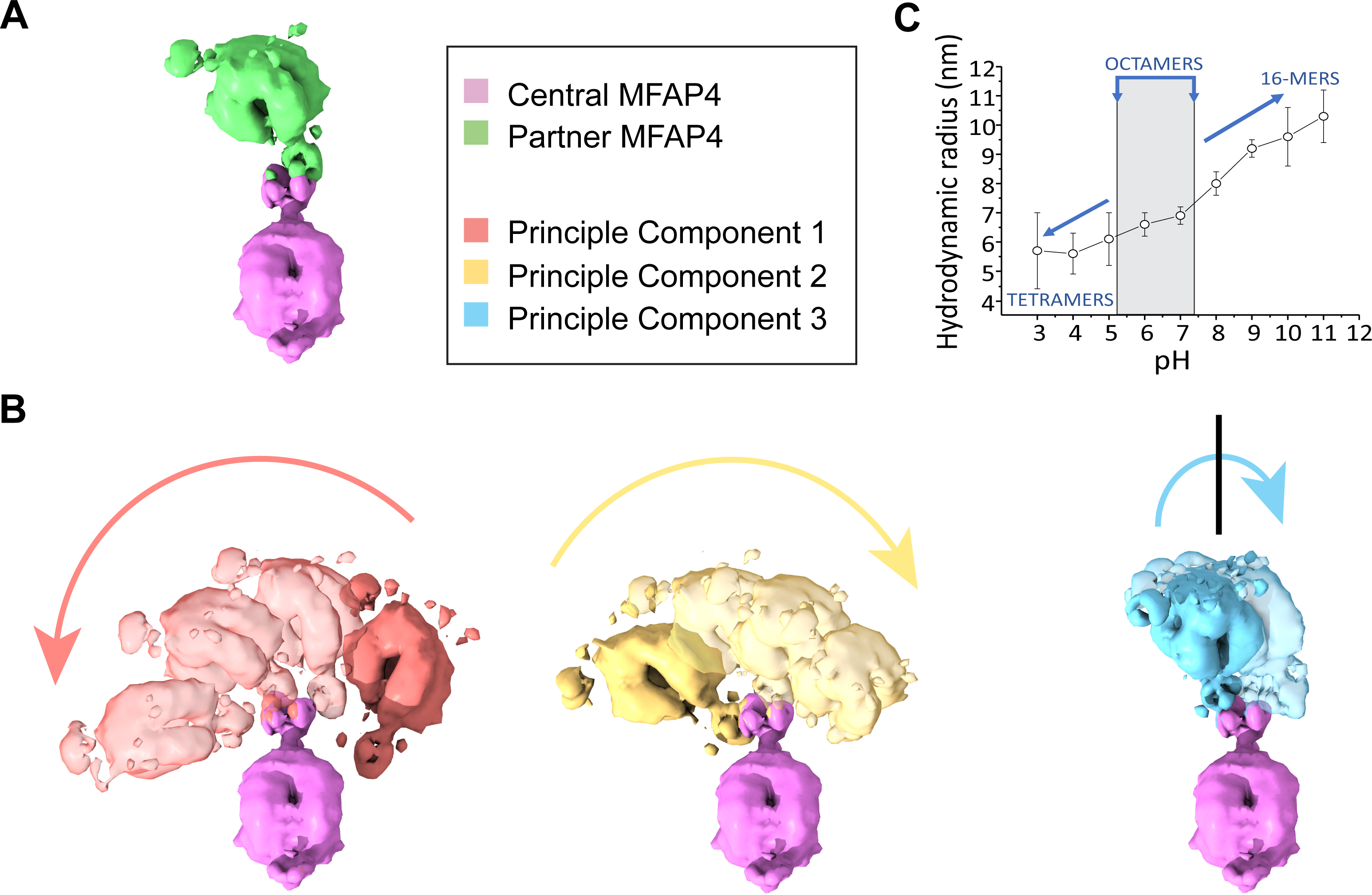
MFAP4 octamer chains. (**A**) Multibody refinement of central MFAP4 (pink) and partner (green) in TBS+Ca^2+^. (**B**) The first, third, eighth and tenth map are shown along the first three principal components of movement. The first (red) and second (yellow) principal components of movement describe a swinging movement of the partner relative to the central MFAP4 from the left to right or right to left, respectively. The third (blue) principal of movement describes a twisting movement of the partner relative to the central octamer. (**C**) Hydrodynamic radii of MFAP4 at pH 3-11 determined by DLS. Octamers are mainly observed at pH 6-7.

## Discussion

MFAP4 is an extracellular protein that has been associated with elastic fibre formation and several diseases with defects in elastic fibres, including Marfan syndrome and chronic obstructive pulmonary disease ^1,3,18^. To advance knowledge of how MFAP4 structurally and functionally relates to elastogenesis and the underpinning diseases, we determined the structure of human MFAP4 by cryo-EM and revealed important structure-function relationships in the context of multimerisation, calcium binding, ligand interaction, glycosylation, and cell interaction.

Previous studies have shown that MFAP4 and its bovine homolog MAGP-36 exists as disulfide-bonded dimers under denaturing conditions ^10,27,35^. The existing paradigm is that these dimers assemble into higher order multimers, albeit there is discrepancy on the type of assembled multimers, i.e. hexamers, octamers, or dodecamers ^10,27^. Here we provide evidence that C34 is sufficient and necessary for intermolecular disulfide bond formation of MFAP4 using the MFAP*_C34S_* mutant. Surprisingly, however, we found that the C34-mediated intermolecular disulfide bond is not required for higher order assembly. For wild type MFAP4 it is presently not known whether the intermolecular disulfide bond forms within the secretory pathway or possibly later after secretion. Our data favour a role of the C34-mediated intermolecular disulfide bond in stabilising higher order assembly as the hydrodynamic radius of the assembled complex decreases more for MFAP*_C34S_* than for MFAP4 with increasing NaCl concentration.

The 3.55 Å resolution cryo-EM structure of human MFAP4 in the presence of Ca^2+^ unequivocally demonstrates an octamer structure and thus solves the existing discrepancy in the field. Octamer formation of MFAP4 contrasts with the ability of other related FReD-containing proteins to multimerise. FIBCD1 forms tetramers (4M7H) ^31^, ficollin-2 forms trimers (2J3G) ^39^, and the FReD of angiopoietin-1 (Ang1) forms dimers (4EPU). A unique conformation of loop A182-A206 in MFAP4 which distinguishes it from other FReD-containing proteins likely explains these differences as outlined below.

In the absence of Ca^2+^, MFAP4 adopts a tetrameric conformation, which represents the identical top or bottom halves of the octamers. Each tetramer contains two sets of protomers linked as dimers by intermolecular disulfide bonds between C34. The fact that after removal of Ca^2+^ tetramers remain stable, support a model of assembly where two MFAP4 protomers associate through intra-tetrameric interactions, predominantly through salt-bridge and hydrogen bonds. The data suggest that Ca^2+^-binding is necessary for the conformation of loop A182-A206 and is stabilised by Ca^2+^-binding to S186, T187, D191, and Q192. This conformation presumably is necessary for inter-tetrameric interactions which are predominantly non-polar, non-bonded interchain interactions between V96-F93, L202-L202, S204-L202 apart from a single polar non-bonded interchain interaction between N231-A200.

In the presence of Ca^2+^ we also observed formation of concatenated octamers forming a chain-like assembly with an approximate distance of 16 nm between octamers. This is likely based on the very high self-assembly affinity of octamers in the low nanomolar range. The octamers interact in a pole-to-pole arrangement, possibly through their exposed N-termini which are not resolved by the cryo-EM map but must be situated near the top and bottom poles of MFAP4 octamers. This view is also supported by the fact that the tetramers observed in the absence of Ca^2+^ never form any chain-like conformation. The preferential orientation of tetramers likely discourages interaction between the N-termini emanating from the centre of tetramers, perpendicular to the monodispersed single layer of vitrified MFAP4 without Ca^2+^. If two tetramers are still able to interact with each other via their poles, it would result in an octamer that cannot further assemble, because of the lack of required Ca^2+^ for the inter-tetrameric interactions. Interestingly, the proposed pole interaction sites for chain formation are situated close to the RGD cell binding site close to the N-terminus, which interacts with integrin αvβ3 and possibly αvβ5 of elastogenic cells, facilitating cell migration and proliferation ^22,30^. MFAP4*_C34S_* interacted somewhat stronger with cells than the wild type intermolecular disulfide-bonded MFAP4. This observation can be readily explained by the structural model that predicts more flexibility of the N-terminus if the C34-mediated disulfide bond is not present. Alternatively, it could reflect a higher propensity of wild type MFAP4 than MFAP4*_C34S_* to form chain-like assemblies potentially obstructing access to the RGD site at the poles. The capacity of MFAP4 or MFAP*_C34S_* cell binding was significantly reduced upon interaction with the N-terminal region of fibrillin-1. Thus, in summary i) the C34-mediated intermolecular disulfide-bond, ii) interaction with fibrillin-1, and iii) possibly chain formation all negatively regulate cell binding to MFAP4. Whether MFAP4 chain formation indeed occurs under physiological conditions and how formation is regulated, however, remains to be investigated.

The interaction survey of MFAP4 with relevant elastogenic proteins revealed Ca^2+^- dependent interactions with fibrillin-1, tropoelastin, fibulin-4, and fibulin-5, with the strongest binding affinity for fibrillin-1 in the low nanomolar range. Binding to fibrillin-1 and tropoelastin was reported previously, albeit MFAP4 interaction with fibrillin-1 was not dependent on Ca^2+^ in that study ^10^. While binding to fibulin-4 and -5 was of moderate affinity, it is possible that MFAP4 is involved in the chaperone function that has been described for fibulin-4 converting folded LTBP4L to an extended conformation which can then align with microfibrils to attract tropoelastin^11^. We identified here that MFAP4 also interacts with an N-terminal region of LTBP4L and LTBP4S, but this interaction is not dependent on Ca^2+^. Together the data support the interpretation that the Ca^2+^-dependent MFAP4 interactions likely require the octameric MFAP4 conformation, whereas the Ca^2+^-independent interactions only require MFAP4 tetramers that are prevalent in the absence of Ca^2+^. MFAP4 octamers could provide unique surfaces for interactions that are unavailable or distorted in the absence of Ca^2+^ as tetramers. Ca^2+^ concentrations in the extracellular matrix are generally assumed to be in equilibrium with circulating blood in the range of 1-2 mM Ca^2+^. Whether or not Ca^2+^ concentrations in specific extracellular spaces, for example in cell invaginations ^40^, can be regulated to a level that would allow MFAP4 tetramers to occur is currently not known. Also, the affinity of MFAP4 for Ca^2+^ that determines the level of Ca^2+^ saturation under physiological conditions remains to be determined. Intriguingly, MFAP4*_C34S_* bound to the tested proteins with nearly identical affinities in the presence of Ca^2+^, with exception of fibulin-5 that did not interact. Another difference in the interaction profile between MFAP4 and MFAP4*_C34S_* was that the latter interacted with LTBP4L and LTBP4S in a Ca^2+^-dependent manner. The data suggest that the interactions of MFAP4 with fibulin-5 and with the LTBP4 isoforms depend on stabilisation of the pole regions by inter-molecular disulfide bonds, but other interactions do not require this stabilisation.

Although FReD family members have different ligand specificities, ligand binding is mediated in part by the S1 pocket, corresponding to aa 187-251 of MFAP4 ^10^. Binding to tropoelastin and collagen type I has previously been investigated by mutating S203 and F241 within the S1 site of MFAP4 ^10^. Both F241A and F241W mutations strongly attenuated or abolished tropoelastin and collagen type I binding. A hydrophobic surface is formed near to the interfaces between tetrameric halves of the octamer by L227, Y229, F241 and Y242; this surface, or at least F241, could represent a potential tropoelastin/collagen type I binding site. Interestingly, the S203Y, but not the S203A mutation also decreased tropoelastin and collagen type I binding ^10^. The cryo-EM map shows that S203 is not exposed in the assembled MFAP4 octamer complex; instead, S203 is tucked within the interface between tetrameric halves of the octamer. We suspect that the S203Y mutation disrupts this interface, possibly by preventing proper folding of the nearby region which is stabilised by a disulfide bond between C199-C212, while S203A is tolerated. Misfolding in this region could potentially prevent C199-C212 disulfide bond formation and/or interfacing between MFAP4 tetramers thus preventing octamer formation. This is in line with tropoelastin requiring the Ca^2+^-saturated MFAP4 octamer for binding but fails to bind to the Ca^2+^ depleted tetramers.

N-linked glycans emanating from N87 and N137 on the surface of MFAP4 octamers suggest that these moieties might influence ligand binding or provide additional ligand binding sites. In the D2 point group symmetric cryo-EM map, both N87 and N137 appear to be simultaneously glycosylated; although, the glycan at N137 is more flexible than that at N87 which has a stronger signal to noise ratio within the cryo-EM map. Interestingly, N87 is closer to the MFAP4 octamer centre of mass and positioned near to the central cavity. It is possible that the glycan at N87 is partially stabilised by this surrounding environment while the glycan bound to N137 can flex within the surrounding solvent. MFAP4 N-linked glycosylation patterns are elevated and more diverse in aneurysmal ascending aortic tissues of patients with Marfan syndrome ^26^. Altered MFAP4 glycosylation could potentially affect binding to elastogenic proteins, including fibrillin-1 or LTBP4 isoforms possibly affecting formation and maintenance of microfibrils and elastic fibres which are compromised in Marfan syndrome. Since the N-linked glycans in MFAP4 are positioned far from F241, it is unlikely that N-glycan alterations would interfere with tropoelastin/collagen type I binding to this site. Given the exposed position on the MFAP4 octamer, the N-linked glycans could also directly promote interactions with elastogenic proteins, possibly altering stability of the elastic fibre system in the ascending aorta. Conversely, C-mannosylation of W235 is likely to play a structural role, given the buried position of W235 within MFAP4. Although we did not observe mannosylation of W235, the target motif (WXXW) is present and previous reports clearly indicate that W235 mannosylation affects MFAP4 secretion ^29^. This difference could arise from the different cell lines used between our study (HEK293) and others (HT1080 human fibrosarcoma). In the event of W235 mannosylation, we would expect a conformational change to accommodate the additional mannose monosaccharide within this buried region.

In the last stages of writing of this manuscript, an X-ray crystal structure of human MFAP4 in complex with a Fab fragment was made available in the Protein Data Bank (PDB:7zmk) by Laursen *et al*. This structure remains undescribed in the literature and is not linked to a manuscript available on a preprint server. Therefore, we cannot discuss differences from a biochemical and functional perspective, but we discuss here the structural differences compared to our cryo-EM structure. Aligned individual monomers exhibit an excellent fit for the core of the molecule, with a root-mean-squared deviation (RMSD) of 0.79Å (198 residue pairs), but a more modest fit for all 220 residues (RMSD: 2.1Å). The largest differences are in the loops F188 - A206 and R210 – N216 that form the tetramer-tetramer contact. When the top tetramer of 7zmk is aligned to the tetramer described in our study, it results in an RMSD of 3.56Å. The bottom tetramers, however, have an RMSD of 19.6Å. This is because the octamer described in our study has a more extended conformation, with a larger central cavity. The 7zmk structure has its lower tetramer rotated so that portions of the lower monomers nestle between two upper monomers. Each monomer then forms contacts with 4 other monomers in the 7zmk structure, instead of only 3 others in our model. In the absence of any information about the biochemical treatment of this MFAP4-Fab complex, we speculate that 7zmk possibly represents an inactive state of MFAP4, since its ligand binding sites are less exposed to solvent.

In conclusion, we present a 3.55 Å resolution structure of human MFAP4 which assembles into octamers and under certain conditions into extended chain-like structures. We clarified the role of Ca^2+^ in the structure and provide extensive structure-function analyses in terms of interaction with elastogenic proteins and cells. This knowledge should provide a solid basis for understanding MFAP4’s role not only in elastogenesis, but also in the various diseases it is associated with.

## Methods

### Protein preparation and purification

For the production of recombinant full length human MFAP4, a double stranded DNA coding for full length MFAP4 including adjacent DNA sequences for cloning was commercially synthesised (Integrated DNA Technologies). The synthetic DNA was cloned into the *NheI* × *NotI* restricted pCEP4 mammalian expression vector (Invitrogen) using the Gibson Assembly Cloning Kit as instructed by the manufacturer (New England BioLabs). The resulting vector was termed pCEP4-MFAP4 coding for the amino acid sequence M^1^KAL…IRRA^255^. An additional hexahistidine tag at the C-terminal end facilitated purification of the recombinant MFAP4. To produce MFAP4*_C34S_*, the codon for C34 in pCEP4-MFAP4 was modified from TGC to TCG using the QuickChange Site Directed Mutagenesis kit as instructed by the manufacturer (Agilent). The entire insert in pCEP4-MFAP4 and the sequence coding for the C34S mutation in pCEP4- MFAP4*_C34S_* was validated by Sanger DNA sequencing. Transfection of 239-EBNA cells, selection, production of conditioned medium containing recombinant MFAP4 or MFAP4*_C34S_*, and purification by nickel chelating chromatography followed previously established procedures ^41,42^. After purification the proteins were dialyzed and stored in 50 mM Tris-HCl, pH 7.4, 150 mM NaCl, 2mM CaCl_2_ (TBS/Ca^2+^).

Production and characterisation of recombinant fibulin-3, fibulin-4 and fibulin-5 ^28^, LTBP4L and LTBP4S ^11^, full length fibrillin-1 ^43^, and the fibrillin-1 subfragments rFBN1-N and rFBN1-C ^44^, and rF1M ^41^, were previously described in detail. Recombinant human tropoelastin was purchased from Elastagen, and purified plasma fibronectin was obtained from Millipore.

### Dynamic light scattering

Particle sizes of MFAP4 and MFAP4*_C34S_* were assessed by dynamic light scattering (DLS) (DynaProMolecular-Sizing Instrument, Protein Solutions, Wyatt Technology). Acquisitions of 5 s or 10 s readings were recorded for each measurement and averaged. Samples of different MFAP4 and MFAP4*_C34S_* preparations in various buffer conditions, either gel filtrated or not, or treated with EDTA were analysed in quadruplicates and the results were averaged. Molecular mass analyses of MFAP4 and MFAP4*_C34S_* from measured hydrodynamic radii were performed applying the globular (compact) model for the conformational protein shape using calibration curves developed from protein standards of known molecular masses provided by the manufacturer (DynaPro).

### Gel filtration chromatography

MFAP4 and MFAP4*_C34S_* were analysed by gel filtration chromatography using an AKTA Avant 25 chromatography system with a 24 mL analytical Superose 12 10/300 column (Cytiva). The proteins (150 µg in 500 µL) were loaded onto the column either in the TBS/Ca^2+^ buffer, or after removal of calcium by 5 mM EDTA incubation for 20 min. The respective elution buffers were 50 mM Tris-HCl, pH 7.4, 500 mM NaCl, 2 mM CaCl_2_ for the calcium-containing condition and 50 mM Tris-HCl, pH 7.4 500 mM NaCl, 5 mM EDTA for the calcium-free condition. The elution volume was collected in 0.3 mL fractions.

### Atomic force microscopy

AFM imaging was performed with a Multimode HR 8 instrument (Bruker) in tapping mode in air. MFAP4 and MFAP4*_C34S_* were absorbed on freshly cleaved Muscovite grade 1 mica surfaces for few minutes and quickly dried with pressurized air. The AFM probes used (TESPA-V2; Bruker) had a nominal tip radius of 2 nm. Quantification of size, shape and abundance of MFAP4 and MFAP4*_C34S_* structures was performed by post-processing analysis of the AFM height images with molecules selected from multiple images and experiments using the instrument software (NanoScope). Image processing included surface flattening as well as adjustments of the Z-axis scale and offset, brightness, and contrast. Particle shape and size quantification was performed to ensure adequate reproducibility of several protein preparations analysed with a minimum of 300 particles per protein preparation and condition. Molecular structures of more complex shapes or aggregates were excluded from the analysis. For some experiments, MFAP4 was mixed with the N-terminal half (rFBN1-N) or full-length recombinant fibrillin-1 (1:1 protein mass concentration; 50 µg/mL of each component) and then imaged by AFM.

### Cryo-electron microscopy, single-particle analysis and atomic modelling

Purified recombinant human MFAP4 was vitrified on CF-2/1-3Cu grids (Protochips, 20 nm carbon, 2.0 μm hole diameter, 1.0 μm hole spacing, 300 mesh copper grid) as 3.5 μL of sample (MFAP4 Stock ID 1289, 0.551 mg/mL, in TBS/Ca^2+^) following 1.5 s of blotting with blot-force 10 using a Vitrobot at 4°C and 90% humidity. Images were collected on a FEI Krios operating at 300 kV at a magnification of 105,000 with beam-tilt using SerialEM. Images were recorded on a Gatan K3 camera with pixel size of 0.855 Å/px. Images were dose-fractionated as 40 frames over a total exposure time of 3.356 s with 0.0839 s per frame and a frame dose of 2.00 electrons per Å^2^. The defocus range was -1.0 to -2.5 μm under focus. 6,751 images were collected.

The data was motion corrected and dose-weighted within RELION4-beta ^45^. Contrast transfer function (CTF) parameter estimation was carried out on the non-dose-weighted, motion-corrected image stacks with CTFFIND4 ^46^. MFAP4 octamers were manually picked for the initial 2D model used for template matching and automatic picked with Topaz within RELION4-beta ^45^. An initial number of 878,597 putative particles were identified within 6,714 micrographs. After 2D classification 458,868 particles were used for initial 3D reconstruction. After 3D refinement and further classification, 455,190 particles contributed to a 3.55 Å resolution structure of the entire MFAP4 octamer, reconstructed with D2 symmetry. Resolution was estimated from the Fourier shell correlation (FSC) at 0.143 from a masked reference with Phenix version 1.20.1-4487_34_.

A starting model of human MFAP4 was predicted with AlphaFold Colab (https://colab.research.google.com/github/deepmind/alphafold/blob/main/notebooks/AlphaFold.i <u>pynb</u>), a simplified version of AlphaFold v2.3.1 ^47^. This starting model was initially docked within the presented cryo-EM map using rigid body fitting in Chimera ^48^. This predicted model was refined manually in Coot (v0.9.8.1) ^33^, then automatically refined within Phenix (v1.20.1-4487) ^34^. Iterative manual and automatic refinement steps were performed with Coot and Phenix to obtain the atomic model. A map-to-model fixed radius cross-correlation score was calculated according to Jiang and Brünger, 1994, with Phenix during automatic map refinement ^49^.

### Protein and cell binding assays

Surface plasmon resonance (SPR) spectroscopy was used to study binding characteristics and kinetics of MFAP4 and MFAP4*_C34S_* interaction with itself and with elastic fibre associated proteins referenced above (fibrillin-1 fragments rFBN1-N (N-terminal half), rFBN1-C (C-terminal half), rF1M (centre region), tropoelastin, LTBP4L (long isoform), LTBP4S (short isoform), LTBP4L/S N-terminal and C-terminal halves, fibulin-3, fibulin-4, fibulin-4, and fibronectin). The experiments were performed with a Biacore X instrument using CM5 sensor chips (Cytiva). Ligand protein MFAP4 or MFAP4*_C34S_* was covalently immobilized onto one of the two channels of the sensor chip by amine coupling using the standard protocol supplied by the manufacturer with immobilisation levels in the range of 300-500 resonance units (RU). The second channel of the sensor chip was left blank (no protein coated) which served as control for analyte binding specificity after inactivation of the reactive amine groups. Kinetic analyses were performed in running buffer (TBS/2mM CaCl_2_) for interaction in the Ca^2+^-form, and in TBS buffer for interaction without Ca^2+^. For Ca-free interactions, some samples were treated with 5 mM EDTA for 20 min followed by dialyses against TBS to remove Ca^2+^. Binding experiments were performed by injecting 30 µL of analyte (0-100 µg/mL) over the sensor surface at 10 µL/min flow rate in TBS/2mM CaCl_2_ or TBS. Molecular association was monitored for 180 s followed by dissociation for 600 s under identical flow conditions. The association-dissociation sensorgrams were fitted using the BIAevaluation software (Cytiva). A 1:1 Langmuir binding model was chosen for fitting and the equilibrium dissociation constant (K*_D_*) characteristic for each interaction was determined. The molecular masses of the analyte proteins used for the calculation of the molar concentrations were calculated from the protein amino acid sequence (theoretical) for monomers as well as octamers/tetramers as applicable. Some sensorgrams were corrected for bulk shift effects occurring occasionally due to sharp refractive index changes at the beginning and the end of analyte injection.

Cell attachment was tested using primary human dermal fibroblasts (local ethics board approval PED-06-054) following established procedures ^50^. Briefly, MFAP4, MFAP4*_C34S_*, fibrillin-1 fragment rFBN1-N (N-terminal half of fibrillin-1), and plasma fibronectin were coated at 50 µg/mL in TBS at 4°C overnight onto 96-well Maxisorp plates (Nunc). In some cases, 50 µg/mL MFAP4 or MFAP4*_C34S_* were preincubated with 50 µg/mL rFBN1-N (N-terminal half of fibrillin-1 ending prior to the RGD-containing TB4 domain ^44^) for 30 min at 22°C, or alternatively, first coated onto the wells at 50 µg/mL and then additionally incubated for 1 h with rFBN1-N (50 µg/mL). Wells were washed with TBS and blocked with 10 mg/mL denatured bovine serum albumin. Near-confluent layers of fibroblasts were briefly trypsinized (3 min), counted and seeded at 25,000 cells/well in 6 technical replicates for 1 h at 37°C. The wells were extensively washed, fixed with 5% (w/v) glutaraldehyde in PBS (20 min at 22°C), washed again and stained with 0.1% (w/v) crystal violet (Sigma Aldrich). After extensive washing with TBS, either the dye was solubilized with 10% (v/v) acetic acid and the absorbance was measured at 570 nm in a spectrophotometer microplate reader (Beckman Coulter, Model AD340), or alternatively, the stained wells were photographed and the optical density was quantified by ImageJ ^51^. Both methods produced identical results. Background values of wells not coated with proteins but including cells were subtracted from the experimental values.

## Acknowledgements

This study was supported by the Natural Sciences and Engineering Research Council of Canada (RGPIN-2022-05045 to DPR and RGPIN-2020-04837 to MS), the Genetic Aortic Disorders Association Canada (DPR), the Canadian Institutes of Health Research (PJT-186194 to MS, Postdoctoral Fellowship MFE-187851 to MRW).

## References

1. Kobayashi, R. et al. Isolation and characterization of a new 36-kDa microfibril-associated glycoprotein from porcine aorta. Journal of Biological Chemistry 264, 17437–17444 (1989).

2. Wulf-Johansson, H. et al. Localization of microfibrillar-associated protein 4 (MFAP4) in human tissues: clinical evaluation of serum MFAP4 and its association with various cardiovascular conditions. PloS One 8, e82243 (2013).

3. Kanaan, R., Medlej-Hashim, M., Jounblat, R., Pilecki, B. & Sorensen, G.L. Microfibrillar-associated protein 4 in health and disease. Matrix Biology 111, 1–25 (2022).

4. Sakai, L.Y., Keene, D.R. & Engvall, E. Fibrillin, a new 350-kD glycoprotein, is a component of extracellular microfibrils. Journal of Cell Biology 103, 2499–2509 (1986).

5. Sabatier, L. et al. Fibrillin assembly requires fibronectin. Molecular Biology of the Cell 20, 846–858 (2009).

6. Kinsey, R. et al. Fibrillin-1 microfibril deposition is dependent on fibronectin assembly. Journal of Cell Science 121, 2696–2704 (2008).

7. Hubmacher, D. et al. Biogenesis of extracellular microfibrils: Multimerization of the fibrillin-1 C-terminus into bead-like structures enables self-assembly. Proceedings of the National Academy of Sciences of the United States of America 105, 6548–6553 (2008).

8. Broekelmann, T.J. et al. Tropoelastin interacts with cell-surface glycosaminoglycans via its COOH-terminal domain. Journal of Biological Chemistry 280, 40939–40947 (2005).

9. Bax, D.V., Rodgers, U.R., Bilek, M.M. & Weiss, A.S. Cell adhesion to tropoelastin is mediated via the C-terminal GRKRK motif and integrin alphaVbeta3. Journal of Biological Chemistry 284, 28616–28623 (2009).

10. Pilecki, B. et al. Characterization of microfibrillar-associated Protein 4 (MFAP4) as a tropoelastin- and fibrillin-binding protein involved in elastic fiber formation. Journal of Biological Chemistry 291, 1103–1114 (2016).

11. Kumra, H. et al. Fibulin-4 exerts a dual role in LTBP-4L-mediated matrix assembly and function. Proceedings of the National Academy of Sciences of the United States of America 116, 20428–20437 (2019).

12. Mecham, R.P. Elastin in lung development and disease pathogenesis. Matrix Biology 73, 6–20 (2018).

13. Papke, C.L. & Yanagisawa, H. Fibulin-4 and fibulin-5 in elastogenesis and beyond: Insights from mouse and human studies. Matrix Biology 37, 142–149 (2014).

14. Sato, F. et al. Lysyl oxidase enhances the deposition of tropoelastin through the catalysis of tropoelastin molecules on the cell surface. Biological and Pharmaceutical Bulletin 40, 1646–1653 (2017).

15. Godwin, A.R.F. et al. The role of fibrillin and microfibril binding proteins in elastin and elastic fibre assembly. Matrix Biology 84, 17–30 (2019).

16. Schiavinato, A. et al. Targeting of EMILIN-1 and EMILIN-2 to fibrillin microfibrils facilitates their incorporation into the extracellular matrix. Journal of Investigative Dermatology 136, 1150–1160 (2016).

17. Ross, R. & Bornstein, P. The elastic fiber: the separation and partial characterization of its macromolecular components. Journal of Cell Biology 40, 366–381 (1969).

18. Toyoshima, T. et al. Ultrastructural distribution of 36-kD microfibril-associated glycoprotein (MAGP-36) in human and bovine tissues. Journal of Histochemistry and Cytochemistry 47, 1049–1056 (1999).

19. Toyoshima, T. et al. Differential gene expression of 36-kDa microfibril-associated glycoprotein (MAGP-36/MFAP4) in rat organs. Cell and Tissue Research 332, 271–278 (2008).

20. Kasamatsu, S. et al. Essential role of microfibrillar-associated protein 4 in human cutaneous homeostasis and in its photoprotection. Scientific Reports 1, 164 (2011).

21. Holm, A.T. et al. Characterization of spontaneous air space enlargement in mice lacking microfibrillar-associated protein 4. American Journal of Physiology. Lung Cellular and Molecular Physiology 308, L1114–24 (2015).

22. Schlosser, A. et al. MFAP4 promotes vascular smooth muscle migration, proliferation and accelerates neointima formation. Arteriosclerosis Thrombosis and Vascular Biology 36, 122–133 (2016).

23. Johansson, S.L. et al. Microfibrillar-associated protein 4: a potential biomarker of chronic obstructive pulmonary disease. Respiratory Medicine 108, 1336–1344 (2014).

24. Kidholm, C.L. et al. Preliminary analysis of proteome alterations in non-aneurysmal, internal mammary artery tissue from patients with abdominal aortic aneurysms. PloS One 13, e0192957 (2018).

25. Bracht, T. et al. Analysis of disease-associated protein expression using quantitative proteomics-fibulin-5 is expressed in association with hepatic fibrosis. Journal of Proteome Research 14, 2278–2286 (2015).

26. Yin, X. et al. Glycoproteomic analysis of the aortic extracellular matrix in Marfan patients. Arteriosclerosis Thrombosis and Vascular Biology 39, 1859–1873 (2019).

27. Schlosser, A. et al. Microfibril-associated protein 4 binds to surfactant protein A (SP-A) and colocalizes with SP-A in the extracellular matrix of the lung. Scandinavian Journal of Immunology 64, 104–116 (2006).

28. Djokic, J., Fagotto-Kaufmann, C., Bartels, R., Nelea, V. & Reinhardt, D.P. Fibulin-3, -4, and -5 are highly susceptible to proteolysis, interact with cells and heparin, and form multimers. Journal of Biological Chemistry 288, 22821-22835 (2013).

29. Osada, Y. et al. The fibrinogen C-terminal domain is seldom C-mannosylated but its C-mannosylation is important for the secretion of microfibril-associated glycoprotein 4. Biochimica et Biophysica Acta - General Subjects 1864, 129637 (2020).

30. Toyoshima, T., Nishi, N., Kusama, H., Kobayashi, R. & Itano, T. 36-kDa microfibril-associated glycoprotein (MAGP-36) is an elastin-binding protein increased in chick aortae during development and growth. Experimental Cell Research 307, 224–230 (2005).

31. Shrive, A.K. et al. Crystal structure of the tetrameric fibrinogen-like recognition domain of fibrinogen C domain containing 1 (FIBCD1) protein. Journal of Biological Chemistry 289, 2880–2887 (2014).

32. Yu, X. et al. Structural basis for angiopoietin-1-mediated signaling initiation. Proceedings of the National Academy of Sciences of the United States of America 110, 7205–7210 (2013).

33. Emsley, P., Lohkamp, B., Scott, W.G. & Cowtan, K. Features and development of Coot. Acta Crystallographica. Section D: Biological Crystallography 66, 486–501 (2010).

34. Liebschner, D. et al. Macromolecular structure determination using X-rays, neutrons and electrons: recent developments in Phenix. Acta Crystallographica. Section D: Structural Biology 75, 861–877 (2019).

35. Kobayashi, R., Mizutani, A. & Hidaka, H. Isolation and characterization of a 36-kDa microfibril-associated glycoprotein by the newly synthesized isoquinolinesulfonamide affinity chromatography. Biochemical and Biophysical Research Communications 198, 1262–1266 (1994).

36. Yariv, B., et al. Using evolutionary data to make sense of macromolecules with a “face-lifted” ConSurf. in Protein Science Vol. 32 e4582 (2023).

37. Sugar, S. et al. Alterations in protein expression and site-specific N-glycosylation of prostate cancer tissues. Scientific Reports 11, 15886 (2021).

38. Nakane, T. & Scheres, S.H.W. Multi-body refinement of cryo-EM images in RELION. Methods in Molecular Biology 2215, 145–160 (2021).

39. Garlatti, V. et al. Structural insights into the innate immune recognition specificities of L- and H-ficolins. EMBO Journal 26, 623–633 (2007).

40. Canty, E.G. et al. Coalignment of plasma membrane channels and protrusions (fibripositors) specifies the parallelism of tendon. Journal of Cell Biology 165, 553–563 (2004).

41. Zeyer, K.A., Zhang, R.M., Kumra, H., Hassan, A. & Reinhardt, D.P. The fibrillin-1 RGD integrin binding site regulates gene expression and cell function through microRNAs. Journal of Molecular Biology 431, 401–421 (2019).

42. Kaur, J. & Reinhardt, D.P. Immobilized metal affinity chromatography co-purifies TGF-beta1 with histidine-tagged recombinant extracellular proteins. PLoS One 7, e48629 (2012).

43. Lin, G. et al. Homo- and heterotypic fibrillin-1 and -2 interactions constitute the basis for the assembly of microfibrils. Journal of Biological Chemistry 277, 50795–50804 (2002).

44. Jensen, S.A., Reinhardt, D.P., Gibson, M.A. & Weiss, A.S. Protein interaction studies of MAGP-1 with tropoelastin and fibrillin-1. Journal of Biological Chemistry 276, 39661–39666 (2001).

45. Kimanius, D., Dong, L., Sharov, G., Nakane, T. & Scheres, S.H.W. New tools for automated cryo-EM single-particle analysis in RELION-4.0. Biochemical Journal 478, 4169–4185 (2021).

46. Rohou, A. & Grigorieff, N. CTFFIND4: Fast and accurate defocus estimation from electron micrographs. Journal of Structural Biology 192, 216–21 (2015).

47. Jumper, J. et al. Highly accurate protein structure prediction with AlphaFold. Nature 596, 583–589 (2021).

48. Pettersen, E.F. et al. UCSF Chimera--a visualization system for exploratory research and analysis. Journal of Computational Chemistry 25, 1605–1612 (2004).

49. Jiang, J.S. & Brünger, A.T. Protein hydration observed by X-ray diffraction. Solvation properties of penicillopepsin and neuraminidase crystal structures. Journal of Molecular Biology 243, 100–115 (1994).

50. Humphries, M.J. Cell adhesion assays. Methods in Molecular Biology 522, 203–210 (2009).

51. Schneider, C.A., Rasband, W.S. & Eliceiri, K.W. NIH Image to ImageJ: 25 years of image analysis. Nature Methods 9, 671–675 (2012).

