## Supplemental Figures for "Microfibril-associated glycoprotein 4 forms octamers that mediate interactions with elastogenic proteins and cells"

**Supplemental Table 1.** MFAP4 interactions with elastogenic full length and sub-fragment proteins in the presence of $Ca^{2+}$ (octameric form) determined by SPR. This summary table includes in addition to the values shown in Figs. 3 and 4 other tested interactions.

| **Binding Ligand** | **$K_D$ (nM)** | **Binding Strength** | **$Ca^{2+}$-Dependency** |
|---|---|---|---|
| MFAP4 (self-interaction) | 6.2 ± 0.9 | Very strong | Yes |
| rFBN1-N (Fibrillin-1, N-terminal half) | 1.8 ± 0.9 | Very strong | Yes |
| rFBN1-C (Fibrillin-1, C-terminal half) | No binding | No binding | Not applicable |
| rF1M (Fibrillin-1, center region) | 1.2 ± 0.5 | Very strong | Yes |
| Tropoelastin | 45 ± 9 | Strong | Yes |
| LTBP4L | 18 ± 5 | Strong | No |
| LTBP4S | 15 ± 6 | Strong | No |
| LTBP4L (N-terminal half) | 19 ± 6 | Strong | Not determined |
| LTBP4S (N-terminal half) | 16 ± 4 | Strong | Not determined |
| LTBP4L/S (C-terminal half) | No binding | No binding | Not applicable |
| Fibulin-3 | No binding | No binding | Not applicable |
| Fibulin-4 | 110 ± 33 | Moderate | Yes |
| Fibulin-5 | 357 ± 47 | Moderate | Yes |
| Fibronectin | No binding | No binding | Not applicable |

**Supplemental Figure Legends**

**Supplemental Figure 1. MFAP4 multimers dissociate into dimers under denaturing conditions and monomers under reducing conditions.** (**A**) The curve shows theoretical hydrodynamic radii plotted against molecular masses of globular proteins. Measured hydrodynamic radii for MFAP4 and MFAP4*C34S* are shown (dashed lines) in the presence or absence of 1% SDS or 50 mM DTT as indicated. (**B**) DLS-determined hydrodynamic radii of MFAP4 (blue and green) and MFAP4*C34S* (red and orange) under denaturing conditions in the presence of 1% SDS in TBS buffer either with or without reducing DTT (50 mM) as indicated. (**C**) Coomassie-stained SDS gel of MFAP4 with decreasing concentrations of DTT (50-0 mM). The position of MFAP4 monomers and dimers are indicated by black arrowheads.

**Supplemental Figure 2. Resolution and angular distribution of MFAP4 with $Ca^{2+}$.** (**A**) Fourier shell correlation (FSC) of the MFAP4 structure in the presence of $Ca^{2+}$. Resolution was calculated from the correlation between two independently refined halves of the data. Resolution at the 0.143 criterion is 3.55 Å without masking and 3.15 Å with masking. (**B**) A 3D representation of the angular distribution of particles used in the MFAP4 structure with $Ca^{2+}$.

**Supplemental Figure 3. Conservation of $Ca^{2+}$binding sites.** (**A**) Top and side views of the 3.55 Å resolution cryo-EM map of MFAP4 with $Ca^{2+}$ and (**B**) the atomic model colour coded using ConSurf [36]. Colour coding indicates conserved (magenta) to variable (blue) residues. (**C**) Close-up views of $Ca^{2+}$-binding sites highlighted in B show $Ca^{2+}$-binding near to glycosylated N137 (green square) and near to the top/bottom tetramer interface (orange square).

**Supplemental Figure 4. Glycosylation of MFAP4 N87 & N137**. (**A**) Top view of the cryo-EM map of MFAP4 with $Ca^{2+}$ with superimposed ribbon model. Two disulfides linked protomers are depicted with surface rendering. (**B**) Sideview with surface rendering and (**C**) without surface rendering. Intermolecular disulfide bonded C34 (yellow), N87 and N137 with NAG glycans (blue), and W235 (green) atoms are shown.

**Supplemental Figure 5. Resolution and angular distribution of MFAP4 without $Ca^{2+}$**. (**A**) Fourier shell correlation (FSC) of the MFAP4 structure without $Ca^{2+}$. Resolution was calculated from the correlation between two independently refined halves of the data. Resolution at the 0.143 criterion is 5.26 Å without masking and 4.46 Å with masking. (**B**) A 3D representation of the angular distribution of particles used in the MFAP4 structure without $Ca^{2+}$.

**Supplemental Movie Legends**

**Movie 1. Cyro-EM map and atomic model of MFAP4 with $Ca^{2+}$.** D2 point group symmetry 3D cryo-EM map of MFAP4 along with superimposed atomic model. Intermolecular disulfide bonded protomer pairs are colour coded accordingly to be the same shade of red or sky-blue. Cysteine residues are coloured yellow. Four N-termini (residues 21-34) of superimposed AlphaFold model is briefly shown aligned to protomers within the presented atomic model; the N-terminal RGD motifs of these AlphaFold models are highlighted (orange). N87 and N137 along with NAG glycans are shown (blue).

**Movie 2. Intra-tetrameric interactions of MFAP4 with $Ca^{2+}$.** Intra-tetrameric interactions within the octamer assembly coloured according to inter-chain distance (dark orange <2.5 Å, light orange <8.5 Å; see colour scale in Fig. 2F/G). Cysteine residues are coloured yellow in all subpanels. Hydrogen bonds are depicted as dashed blue lines. $Ca^{2+}$ ions are depicted as green spheres.

**Movie 3. Inter-tetrameric interactions of MFAP4 with $Ca^{2+}$.** Inter-tetrameric interactions within the octamer assembly coloured according to inter-chain distance (dark orange <2.5 Å, light orange <8.5 Å; see colour scale in Fig. 2F/G). Cysteine residues are coloured yellow in all subpanels. Hydrogen bonds are depicted as dashed blue lines. $Ca^{2+}$ ions are depicted as green spheres.

**Movie 4. Central and partner octamers within MFAP4 chain-like assemblies.** Rotated movie of the central MFAP4 (pink) and partner (green) octamer in TBS/$Ca^{2+}$ following multibody refinement. The first three principal components of movement are also shown, corresponding to the first (red) and second (yellow) swinging movement and the third twisting movement (blue) of the partner relative to the central MFAP4.

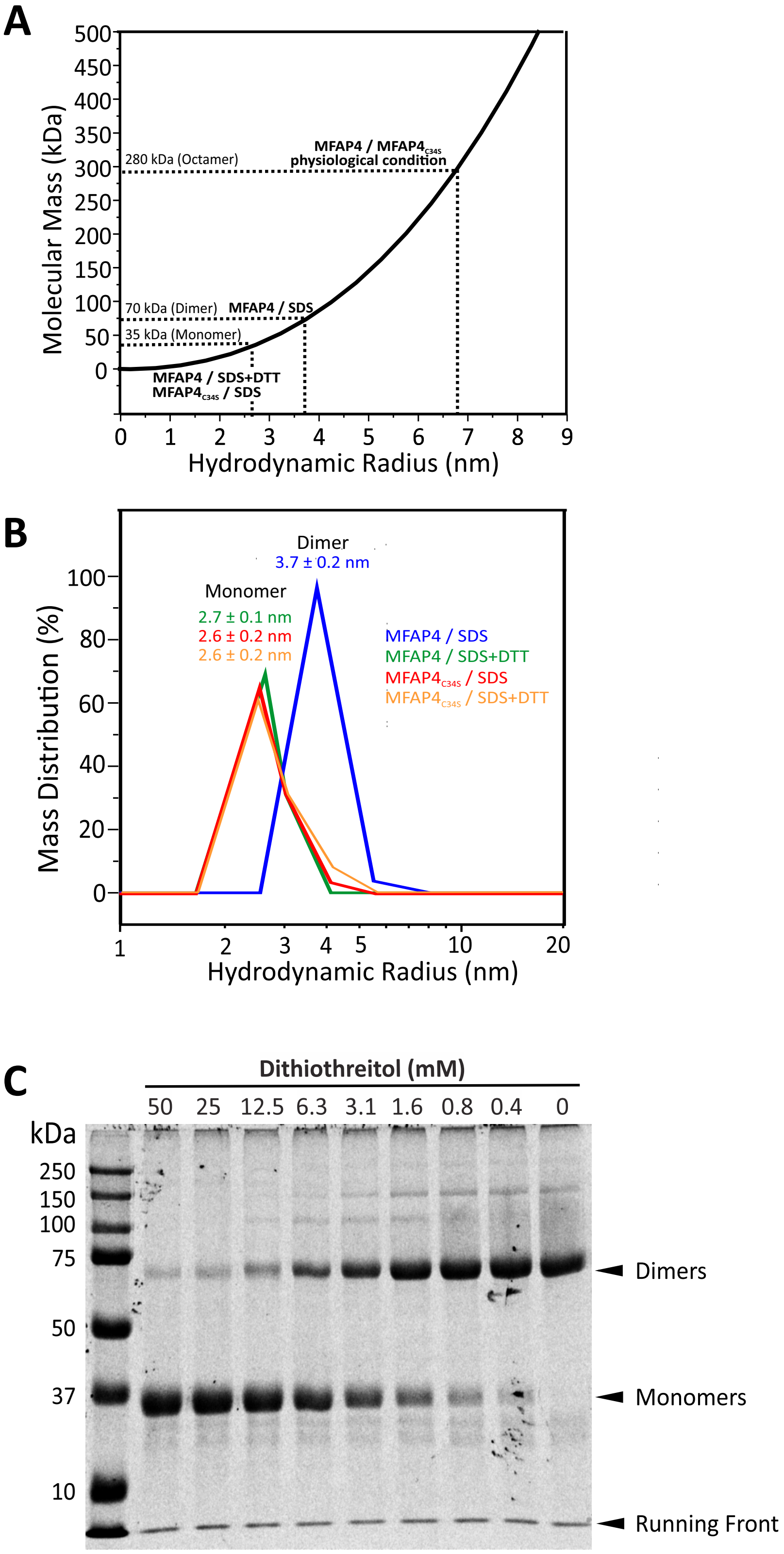
A
Molecular Mass (kDa)
Hydrodynamic Radius (nm)
280 kDa (Octamer)
MFAP4 / MFAP4C34S
physiological condition
70 kDa (Dimer)
MFAP4 / SDS
35 kDa (Monomer)
MFAP4 / SDS+DTT
MFAP4C34S / SDS
B
Mass Distribution (%)
Hydrodynamic Radius (nm)
Dimer
3.7 ± 0.2 nm
Monomer
2.7 ± 0.1 nm
2.6 ± 0.2 nm
2.6 ± 0.2 nm
MFAP4 / SDS
MFAP4 / SDS+DTT
MFAP4C34S / SDS
MFAP4C34S / SDS+DTT
C
Dithiothreitol (mM)
50 25 12.5 6.3 3.1 1.6 0.8 0.4 0
kDa
250
150
100
75
50
37
10
Dimers
Monomers
Running Front

Suppl. Figure 2

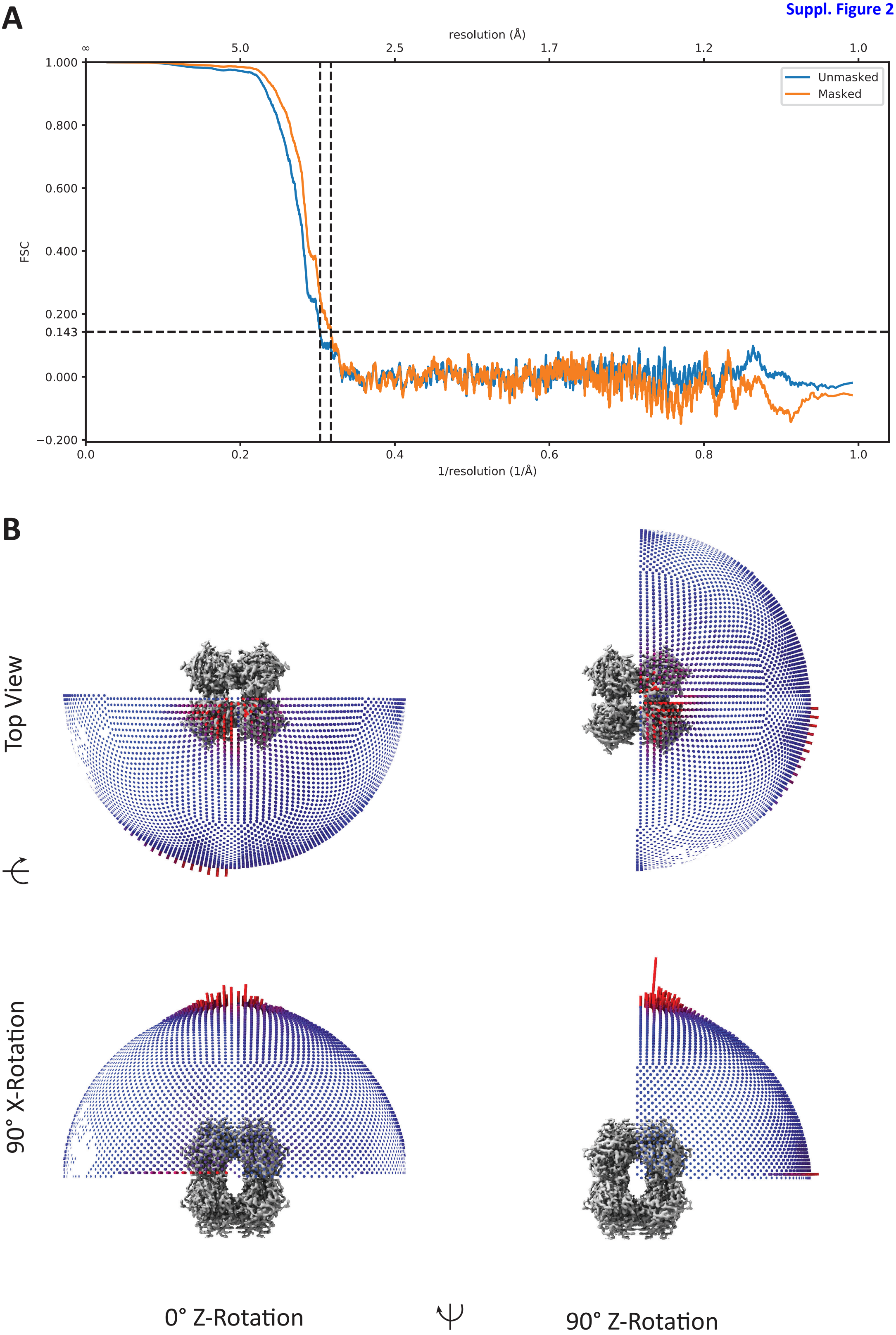

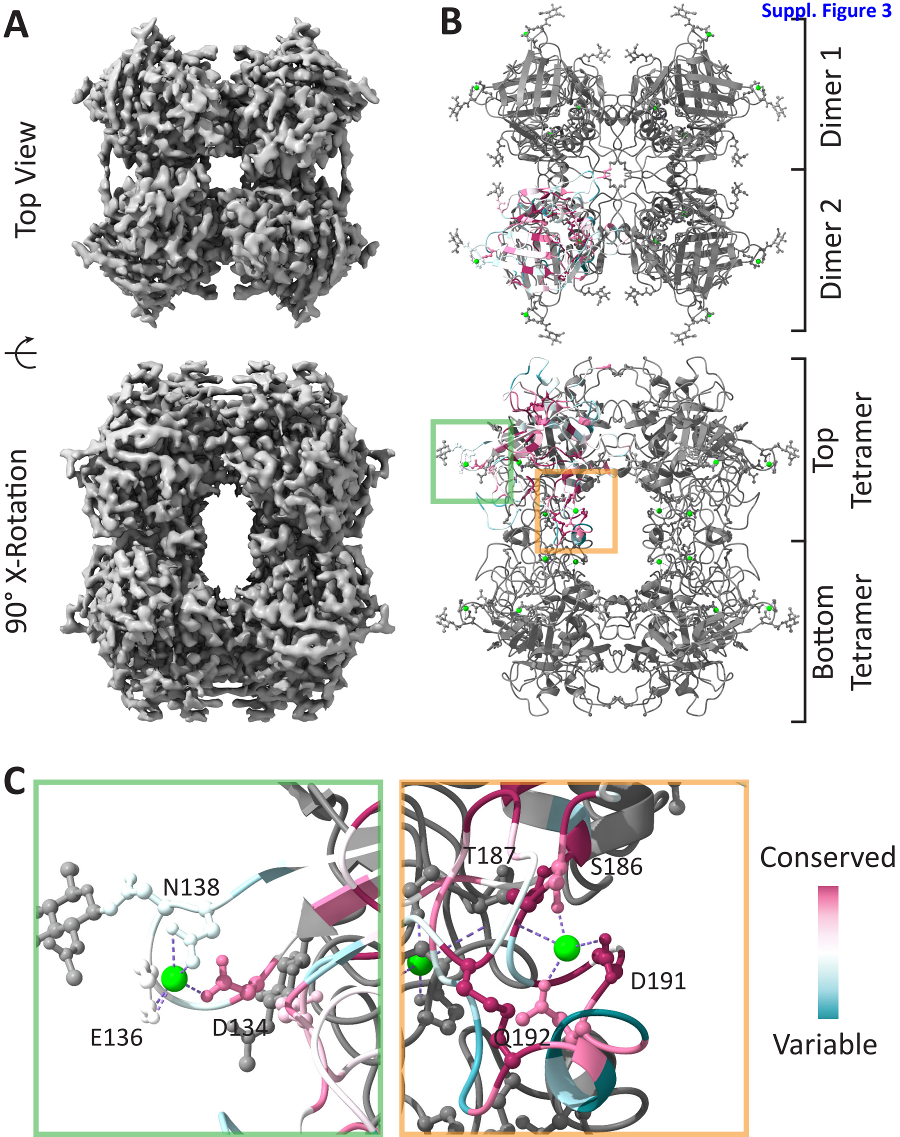
A
B
Top View
90° X-Rotation
Dimer 1
Dimer 2
Top Tetramer
Bottom Tetramer
C
N138
E136
D134
T187
S186
D191
Q192
Conserved
Variable

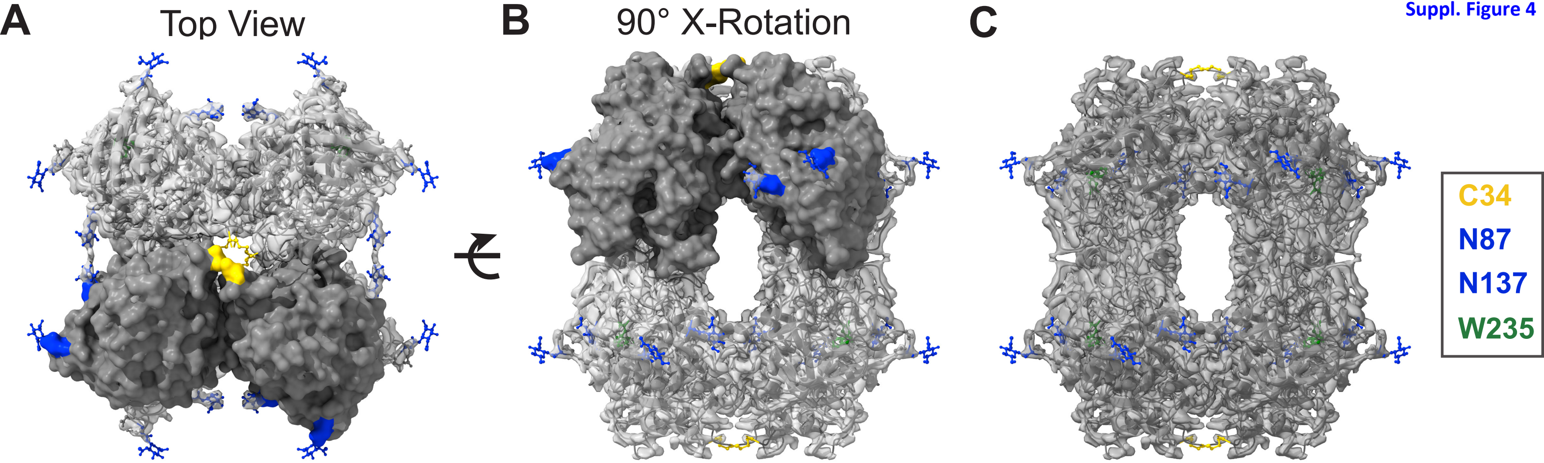
A
Top View
B
90° X-Rotation
C
C34
N87
N137
W235

**Suppl. Figure 5**

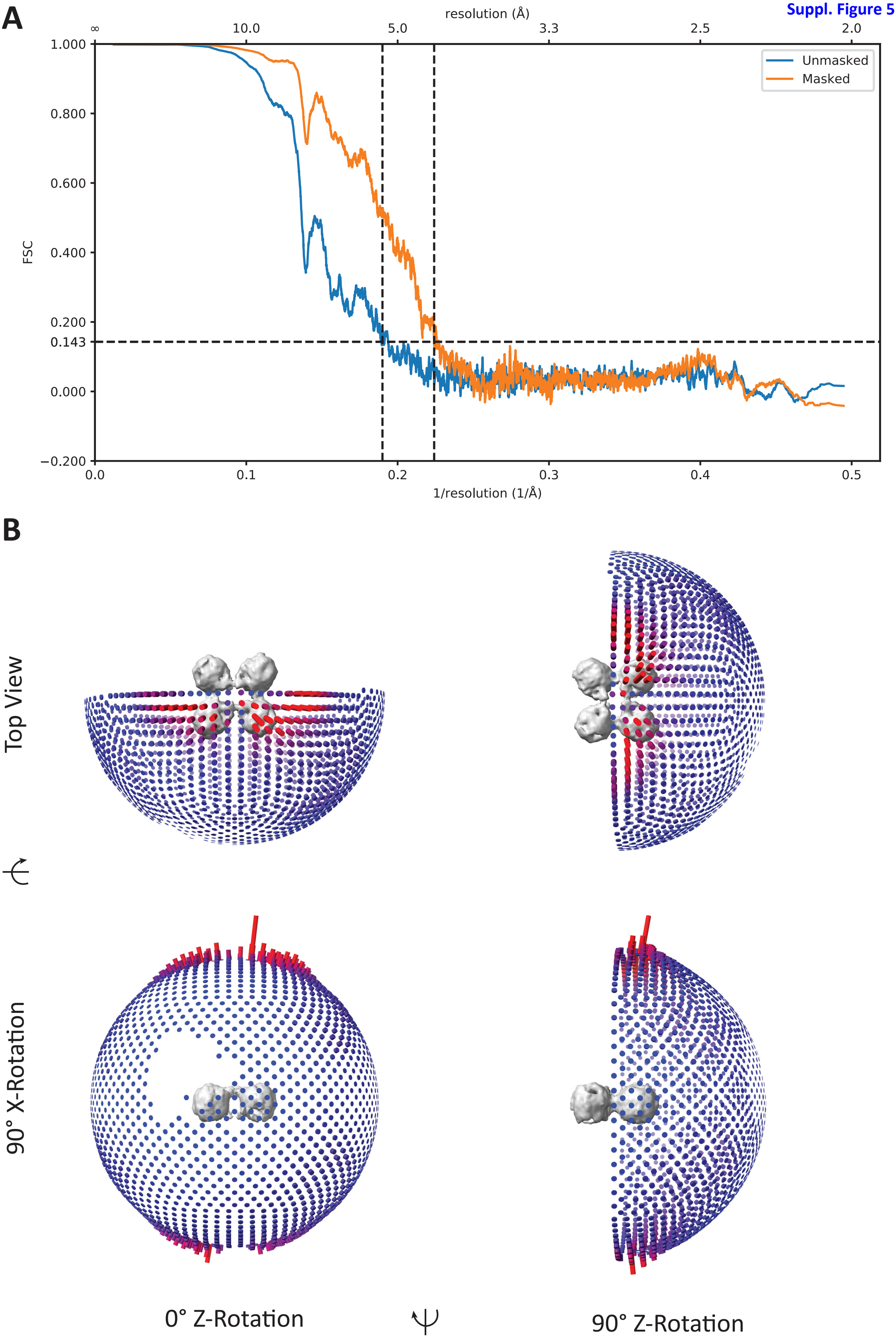
